# High resolution mapping of the breast cancer tumor microenvironment using integrated single cell, spatial and in situ analysis of FFPE tissue

**DOI:** 10.1101/2022.10.06.510405

**Authors:** Amanda Janesick, Robert Shelansky, Andrew D. Gottscho, Florian Wagner, Morgane Rouault, Ghezal Beliakoff, Michelli Faria de Oliveira, Andrew Kohlway, Jawad Abousoud, Carolyn A. Morrison, Tingsheng Yu Drennon, Seayar H. Mohabbat, Stephen R. Williams, 10x Development Teams, Sarah E.B. Taylor

## Abstract

Single cell and spatial technologies that profile gene expression across a whole tissue are revolutionizing the resolution of molecular states in clinical tissue samples. Commercially available methods that characterize either single cell or spatial gene expression are currently limited by low sample throughput and/or gene plexy, lack of on-instrument analysis, and the destruction of histological features and epitopes during the workflow. Here, we analyzed large, serial formalin-fixed, paraffin-embedded (FFPE) human breast cancer sections using a novel FFPE-compatible single cell gene expression workflow (Chromium Fixed RNA Profiling; scFFPE-seq), spatial transcriptomics (Visium CytAssist), and automated microscopy-based in situ technology using a 313-plex gene panel (Xenium In Situ). Whole transcriptome profiling of the FFPE tissue using scFFPE-seq and Visium facilitated the identification of 17 different cell types. Xenium allowed us to spatially resolve these cell types and their gene expression profiles with single cell resolution. Due to the non-destructive nature of the Xenium workflow, we were able to perform H&E staining and immunofluorescence on the same section post-processing which allowed us to spatially register protein, histological, and RNA data together into a single image. Integration of data from Chromium scFFPE-seq, Visium, and Xenium across serial sections allowed us to do extensive benchmarking of sensitivity and specificity between the technologies. Furthermore, data integration inspired the interrogation of three molecularly distinct tumor subtypes (low-grade and high-grade ductal carcinoma in situ (DCIS), and invasive carcinoma). We used Xenium to characterize cellular composition and differentially expressed genes within these subtypes. This analysis allowed us to draw biological insights about DCIS progression to infiltrating carcinoma, as the myoepithelial layer degrades and tumor cells invade the surrounding stroma. Xenium also allowed us to further predict the hormone receptor status of tumor subtypes, including a small 0.1 mm^2^ DCIS region that was triple positive for *ESR1* (estrogen receptor), *PGR* (progesterone receptor), and *ERBB2* (human epidermal growth factor receptor 2, a.k.a. HER2) RNA. In order to derive whole transcriptome information from these cells, we used Xenium data to interpolate the cell composition of Visium spots, and used Visium whole transcriptome information to discover new biomarkers of breast tumor subtypes. We demonstrate that scFFPE-seq, Visium, and Xenium independently provide information about molecular signatures relevant to understanding cancer heterogeneity. However, it is the integration of these technologies that leads to even deeper insights, ushering in discoveries that will progress oncology research and the development of diagnostics and therapeutics.

## Introduction

High-throughput methods in single cell genomics have made it possible to cluster thousands to millions of cells from a single experiment into distinct types based on whole transcriptome gene expression and cell surface protein data, sparking ambitious collaborations to profile every cell type in the human body (Zheng et al. 2017, Regev et al. 2017, He et al. 2020, Karlsson et al. 2021, Eraslan et al. 2022). Meanwhile, advances in spatial transcriptomics have introduced unbiased gene expression analysis with spatial context for tissue sections, combining genomics, imaging, and tissue pathology (Rao et al. 2020, Maynard et al. 2021). The 10x Genomics Chromium (single cell) and Visium (spatial) platforms are complementary in that Chromium data have single cell resolution, but lack spatial context, while Visium data have spatial context, but may require integration with single cell data to infer detailed information about cell type composition. Both technologies require destruction of tissues during the experimental workflow, necessitating that hematoxylin and eosin (H&E) and immunofluorescence (IF) staining be performed prior or on serial sections. Although there has been progress in integrating these datasets computationally downstream through transcript distribution prediction and cell type deconvolution (Li et al. 2022), high resolution cell-cell and ligand-receptor interactions that comprise intercellular communication are lacking, as is the definitive assignment of transcripts to a particular cell with spatial context at high gene plexy. An ideal solution would provide high-plex, high throughput, multi-modal read outs with spatial context and subcellular resolution, without compromising tissue integrity, and be compatible with both fresh frozen (FF) and formalin-fixed paraffin embedded (FFPE) tissues.

Here, we introduce the novel Xenium In Situ Technology, that offers the aforementioned features, along with a large imageable area and integration of gene expression with histological images (H&E and IF staining) in the same tissue section. The initial commercial Xenium kits will support up to 400 gene plexy with up to 100 add-on custom RNA targets that can be spiked into pre-designed panels, and a throughput of up to ~2.8 cm^2^ of imageable area per slide (up to ~17 cm^2^ per week). The platform is built to support an even higher throughput, gene plexy over 1000, and protein and RNA measurements on the same section. We also introduce RNA templated ligation (RTL) technology for the Chromium platform (Fixed RNA Profiling) and apply it to FFPE tissues (scFFPE-seq), unlocking vast biobanks of samples while also improving sensitivity (Vallejo et al. 2022). Using scFFPE-seq, Visium and Xenium on serial sections of a single FFPE-preserved breast cancer tissue block (Fig. 1), we demonstrate how whole transcriptome and targeted in situ data can be integrated to provide highly complementary and additive biological information. scFFPE-seq and Visium allowed us to annotate the cell types in the sample, which was further refined by mapping the transcripts to the Xenium data. Xenium provides subcellular spatial resolution, which is particularly suited for studying tumor invasion in ductal carcinoma in situ (DCIS), due to its high molecular complexity and close proximity of different cell types. Furthermore, using Xenium, we identified a cell type positive for the RNA of three breast cancer classifying receptors (estrogen, progesterone, and HER2) that the other technologies did not detect. Integration of Visium and Xenium data allowed us to derive high resolution spatially resolved whole transcriptome information for this group of cells, revealing differentially expressed genes associated with the triple-positive tumor region. These findings highlight the synergistic relationship between single cell and spatial data, conspiring to provide a deeper understanding of the complex and diverse network of cells within the tumor microenvironment.

**Figure 1.**
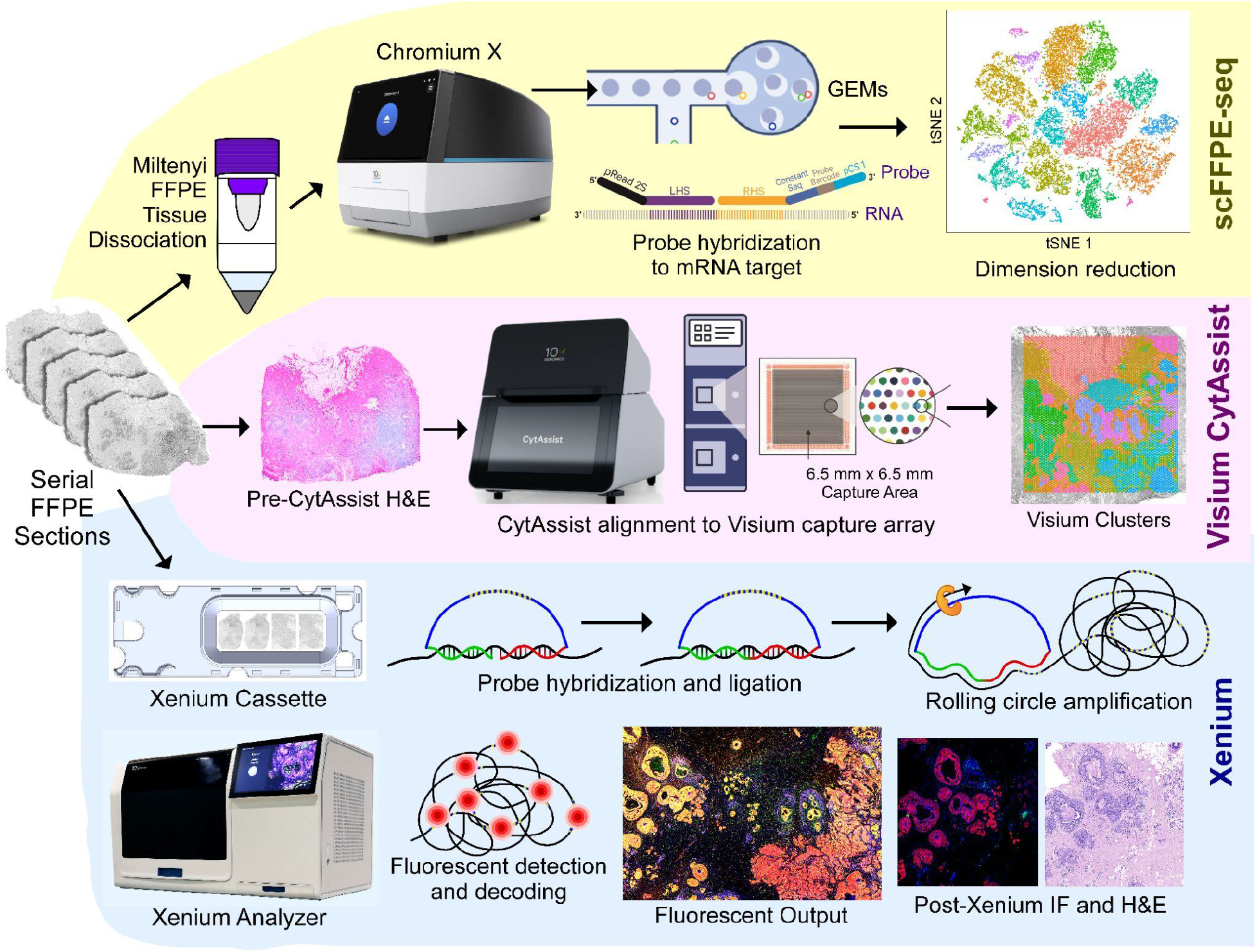
Experimental design. A single FFPE tissue block was analyzed with a trio of complementary technologies. Top: the Chromium Fixed RNA Profiling workflow (scFFPE-seq) with the Miltenyi FFPE Tissue Dissociation protocol. Middle: Visium CytAssist enabled whole transcriptome analysis with spatial context, and was readily integrated with single cell data from serially adjacent FFPE tissue sections. Bottom: The novel Xenium In Situ platform used a microscopy based read-out. A 5 μm tissue section was sectioned onto a Xenium slide, followed by hybridization and ligation of specific DNA probes to target mRNA, followed by rolling circle amplification. The slide was placed in the Xenium Analyzer instrument for multiple cycles of fluorescent probe hybridization and imaging. Each gene has a unique optical signature, facilitating decoding of the target gene, from which a spatial transcriptomic map was constructed across the entire tissue section. The Xenium data could be easily registered with post-Xenium IF / H&E images (as the workflow is non-destructive to the tissue) and integrated with scFFPE-seq and Visium data.

## Results

### Single cell FFPE and Visium data collectively provide whole transcriptome information with spatial context from human breast cancer FFPE tissue

Breast cancer is a complex disease of multiple pathologies: each tumor subtype has unique features and significant cellular and molecular heterogeneity. To better understand tumorigenesis and the cancer ecosystem, it is necessary to dissect cellular components and molecular profiles within the spatial context of the tumor landscape. Using discovery-based technologies, we characterized a breast cancer sample with both single cell and spatial whole transcriptome data. First, we generated Chromium scFFPE-seq data from 2 x 25 μm FFPE curls (see Methods) of a breast cancer block (Stage II-B, ER+/PR−/HER2+) that were adjacent to the tissue sections used for Visium and Xenium workflows. Analysis of the scFFPE-seq data yielded 17 well-segregated clusters based on unsupervised clustering analysis, with a median of 1,480 genes identified per cell.

Next, we generated Visium whole transcriptome data by collecting 5 μm tissue sections adjacent to those used for scFFPE-seq. Sections were H&E stained prior to imaging, followed by Visium CytAssist library preparation and sequencing. The CytAssist instrument facilitates the transfer of analytes from standard glass slides to Visium slides, and uses an updated human probe set, identical to those in the Chromium scFFPE-seq product (18,536 genes targeted by 54,018 probes in the Visium CytAssist probe set vs. 17,943 genes targeted by 18,630 probes in the Visium for FFPE v1 probe set). Dimensionality reduction of the Visium data yielded 17 spatial clusters (coincidently, the same number of clusters as the scFFPE-seq data), with a median of 5,712 genes identified per spot.

With these two discovery-based datasets in hand, and the guidance of existing human breast cancer references (Karlsson et al. 2021), we annotated the scFFPE-seq clusters (Fig. 2A) and mapped cell types onto Visium data (Fig. 2B, C) using an iterative process. Ten Visium clusters were annotated such that they could be unequivocally assigned to cell types or disease states (Fig. 2B), while the other seven clusters had mixed cell type compositions. Visium pinpointed the spatial location of three tumor domains that were revealed as distinct clusters by scFFPE-seq, including two molecularly distinct types of ductal carcinoma in situ (DCIS), named here DCIS #1 and #2, and invasive tumor (Fig. 2C). The Visium workflow also delineated the general territory of immune and stromal cells and was able to recover transcripts from adipocytes, a delicate cell type that can rupture and/or stick to plastic surfaces during dissociation (Picon-Ruiz et al. 2020, Benitez & Shonida 2020; Fig. 2C).

**Figure 2.**
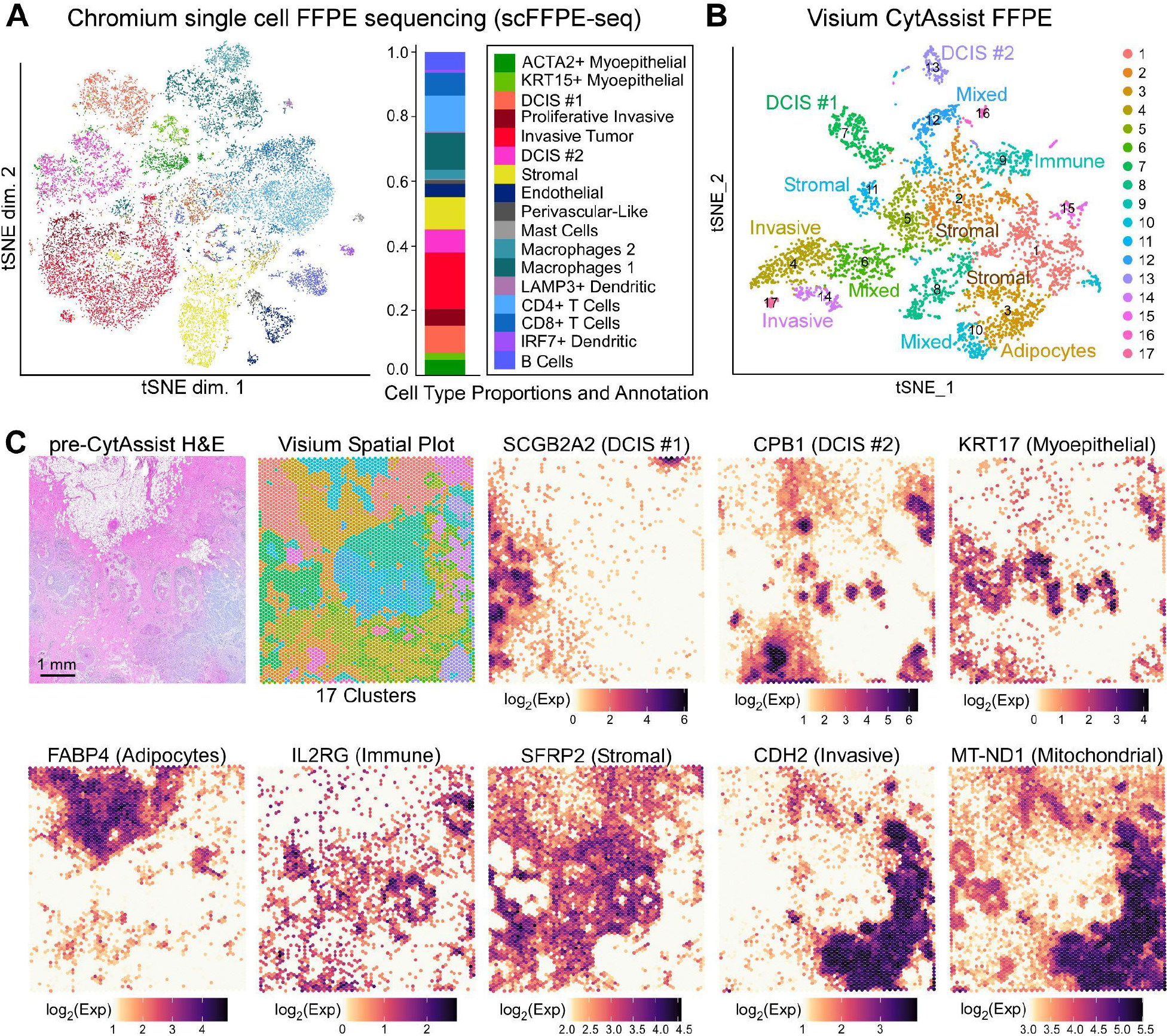
Chromium scFFPE-seq and Visium CytAssist data provide whole transcriptome information with spatial context from FFPE tissues. A human breast cancer sample was obtained as an FFPE block (annotated by pathologist as invasive ductal carcinoma) and processed for single cell analysis and spatial transcriptomics as described in Fig. 1. (A) Dimension reduction of the scFFPE-seq data yielded a t-SNE projection with 17 unsupervised clusters. Each point represents a cell and the colors/labels show annotated cell types (derived from Wu et al. 2021 and Karlsson et al. 2021). (B) t-SNE projection of Visium spots also identifies 17 clusters by unsupervised clustering. Based on differential gene expression analysis, ten clusters could be unequivocally assigned to cell types, while the others were mixtures of cell types. (C) H&E staining conducted pre-CytAssist is shown for reference alongside the spatial distribution of clusters in (B). Cell type-specific marker genes are expressed as log_2_(normalized UMI counts). The Visium data elucidated the spatial location of two molecularly distinct DCIS and invasive subtypes and the general locations of immune, myoepithelial, adipocytes, and stromal cells. Additionally, Visium CytAssist features mitochondrial probes (e.g., MT-ND1), and their spatial distribution correlates with the invasive region of the tissue section.

scFFPE-seq and Visium technologies resolved cellular heterogeneity at single cell level and provided spatial insights, respectively. The integration of scFFPE-seq and Visium data was instrumental to locating cell types and transcripts within the human breast cancer tissue section. However, areas where cell types coexist in close proximity cannot be precisely spatially segregated within the tissue. For example, we observed substantial overlap between DCIS, myoepithelial, immune and stromal markers, and unannotated Visium clusters representing mixtures of cell types. Thus, we next set out to further decipher the cellular composition of the human breast cancer sample in particular, to resolve gene expression within the myoepithelial layer thinly sandwiched between the glandular epithelial cells, the basement membrane and the surrounding stroma.

### Xenium data provide spatially resolved expression of genes at single cell resolution

We next used the Xenium workflow to produce extremely high resolution gene expression data for a targeted panel of genes (Fig. 3). We used 280 genes from the Xenium Human Breast Panel with 33 add-on genes for a total of 313 genes, selected and curated primarily based on single cell atlas data for human breast tissues, including healthy and tumorigenic states (Pal et al. 2021, Bhat-Nakshatri et al. 2021, Karlsson et al. 2021) (Supp. Fig. 1). We visualized the raw fluorescence image after one cycle of decoding, revealing the detailed structure of the tissue with high resolution (Fig. 3A). Further interrogation of the tissue using the Xenium Explorer software allowed us to select relevant genes from the panel to identify stromal, lymphocytes, macrophages, myoepithelial, endothelial, DCIS, and invasive tumor cells (Fig. 3B). We also conducted post-Xenium H&E using standard staining protocols (Fig. 3C), which demonstrates that tissue integrity remains intact even after the full Xenium workflow.

**Figure 3.**
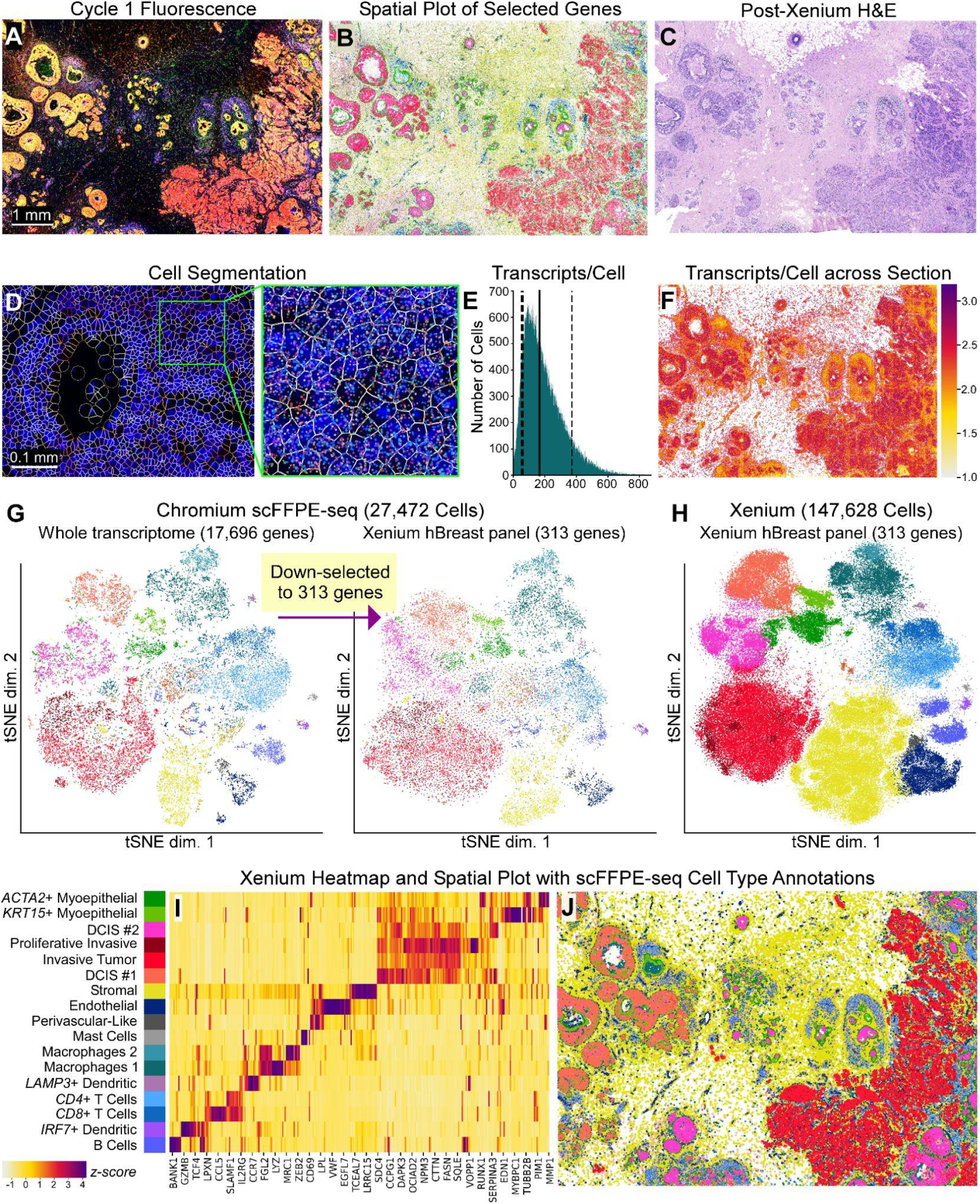
Xenium data provide extremely high resolution single cell information with spatial localization from a targeted panel of genes. (A) Maximum intensity projection of raw fluorescence signal in Cycle 1 from a 5 μm FFPE section. Fifteen of such images (unprojected, original z-stacks), one per cycle, were input into the on-instrument pipeline to decode 313 genes. (B) Selected genes representing major cell types are shown: stromal (*POSTN*), lymphocytes (*IL7R*), macrophages (*ITGAX*), myoepithelial (*ACTA2*, *KRT15*), endothelial (*VWF*), DCIS (*CEACAM6*), and invasive tumor (*FASN*). (C) H&E staining performed post-Xenium workflow, highlighting the minimal impact of the Xenium assay on tissue integrity. (D) Deep learning-based cell segmentation assigns individual transcripts to cells. (E) Histogram showing the distribution of transcripts per cell (Q ≥ 20). Dotted lines: 10th percentile = 61 and 90th percentile = 372 median transcripts per cell. Solid line: 50th percentile = 166 median transcripts per cell. (F) Log_10_(transcripts per cell) across the entire section. (G) t-SNE projection of scFFPE-seq data using all 17,696 genes (left) then down-selected to 313 genes (right). (H) t-SNE projection of labeled Xenium cells. Cell annotation was produced through supervised labeling from scFFPE-seq data. Cells which were not unambiguously identified in the Xenium data (<50% of the nearest neighbors coming from one cell type) were unlabeled (~14% of cells). (I) Heatmap representation of the t-SNE (H) showing the relative expression of genes across different cell types found in the Xenium data. Scale bar is a z-score computed across cell types for each gene, by subtracting the mean and dividing by the standard deviation. See Supp. Fig. 5 for the corresponding scFFPE-seq heatmap. (J) Spatial plot with cell type labels transferred.

### Cell segmentation of Xenium data enables downstream integration and benchmarking with Chromium and Visium data

The gene-cell matrix, or gene-spot matrix, is a standard output format in single cell and spatial transcriptomics that can be input in a variety of community developed tools, including those that integrate different data modalities (e.g. Seurat). In order for Xenium data to be output in this format, it is first necessary to define cell boundaries on the image(s) and then assign transcripts to cells (analogous to the cell-calling stages in single cell transcriptomics). To accomplish this, nuclei were detected from DAPI and expanded outwards until either 15 μm maximum distance was reached, or the boundary of another cell was reached (see Methods). Cell segmentation boundaries can be visualized using the Xenium Explorer software (Fig. 3D), and the on-instrument pipeline outputs Xenium data in standard gene-cell matrix formats, in which transcripts are explicitly assigned to cells. In the section analyzed here, we observed 167,885 total cells, 36,944,521 total transcripts (Q score ≥ 20; see Methods), with a median of 166 transcripts per cell (Fig. 3E, F). When we downsampled the scFFPE-seq data to the 313 genes on the Xenium panel, we observed a median of 34 genes per cell for scFFPE-seq compared to a median of 62 genes per cell in the Xenium data (Supp. Fig. 2A, B). Fifty percent of total transcripts observed contribute to 27 genes (i.e., complexity measurement; Supp. Fig. 2C). Observed counts of negative controls were minimal (Supp. Fig. 2D). Negative control probes accounted for 0.026% of the total counts (Q ≥ 20). Decoding controls accounted for 0.01% of the total counts (Q ≥ 20).

### Chromium scFFPE-seq data confirms that the Xenium gene panel represents major cell types within human breast cancer tissue

To validate our 313-plex human breast Xenium panel, we explored the relative expression of panel genes in expected cell types. The scFFPE-seq and Xenium data were converted to the same gene-cell matrix format for dimension reduction and t-SNE analysis. We transferred supervised scFFPE-seq annotations (see Fig. 2A) to the Xenium data (Supp. Fig. 3); 86% of cells were unambiguously identified as a single cell type in the Xenium data. We filtered the whole transcriptome scFFPE-seq data (17,696 genes; Fig. 2A) to only the 313 genes used in the Xenium human breast panel and found that the same cell type populations were identified (Fig. 3G), confirming that the Xenium human breast panel faithfully captures biological heterogeneity.

The accurate assignment of transcripts to cells allows the same expected cell types to be identified from the Xenium data as from the single cell data, albeit with much higher density due to the greater number of cells analyzed in Xenium (Fig. 3H, I). We mapped the localization of the cell types identified to generate a Xenium spatial plot (Fig. 3J), which can be explored interactively here (also see Data Sharing). Adipocytes were the only cell type not mapped to the Xenium data because scFFPE-seq did not segregate these cells as a distinct cluster based on known adipocyte markers (Supp. Fig. 4A). Xenium, like Visium, successfully identified the location of adipocyte markers, but provided refined resolution where adipocyte transcripts skirt the edge of the cell boundary, since triglycerides fill the majority of the cell (Supp. Fig. 4B-F). Analysis of two serial sections demonstrated the reproducibility of the technology, with the replicates having cell type proportions that were nearly identical and transcript counts that were highly correlated (r^2^ = 0.99) (Supp. Fig. 6).

### Xenium detects RNA transcripts with high sensitivity and specificity

Xenium and scFFPE-seq are new technologies, and therefore, it is prudent to benchmark their sensitivity against each other and relative to existing Chromium technologies that use fresh or frozen (rather than fixed) cells. We quantified sensitivity using median gene expression such that high or low expressors would not bias our measurement. When sequencing depth was kept constant across platforms (~10,000 reads per cell), the median gene sensitivity of scFFPE-seq was higher than the existing 10x single cell platforms (Chromium 5’ Gene Expression (GEX) and 3’ GEX) (Supp. Fig. 7). To benchmark Xenium and scFFPE-seq, we compared the number of transcripts per cell (Xenium) to the number of UMIs per cell (scFFPE-seq), downsampling to the number of genes on the Xenium panel. We found that Xenium is 1.4x more sensitive than scFFPE-seq (Fig. 4A).

**Figure 4.**
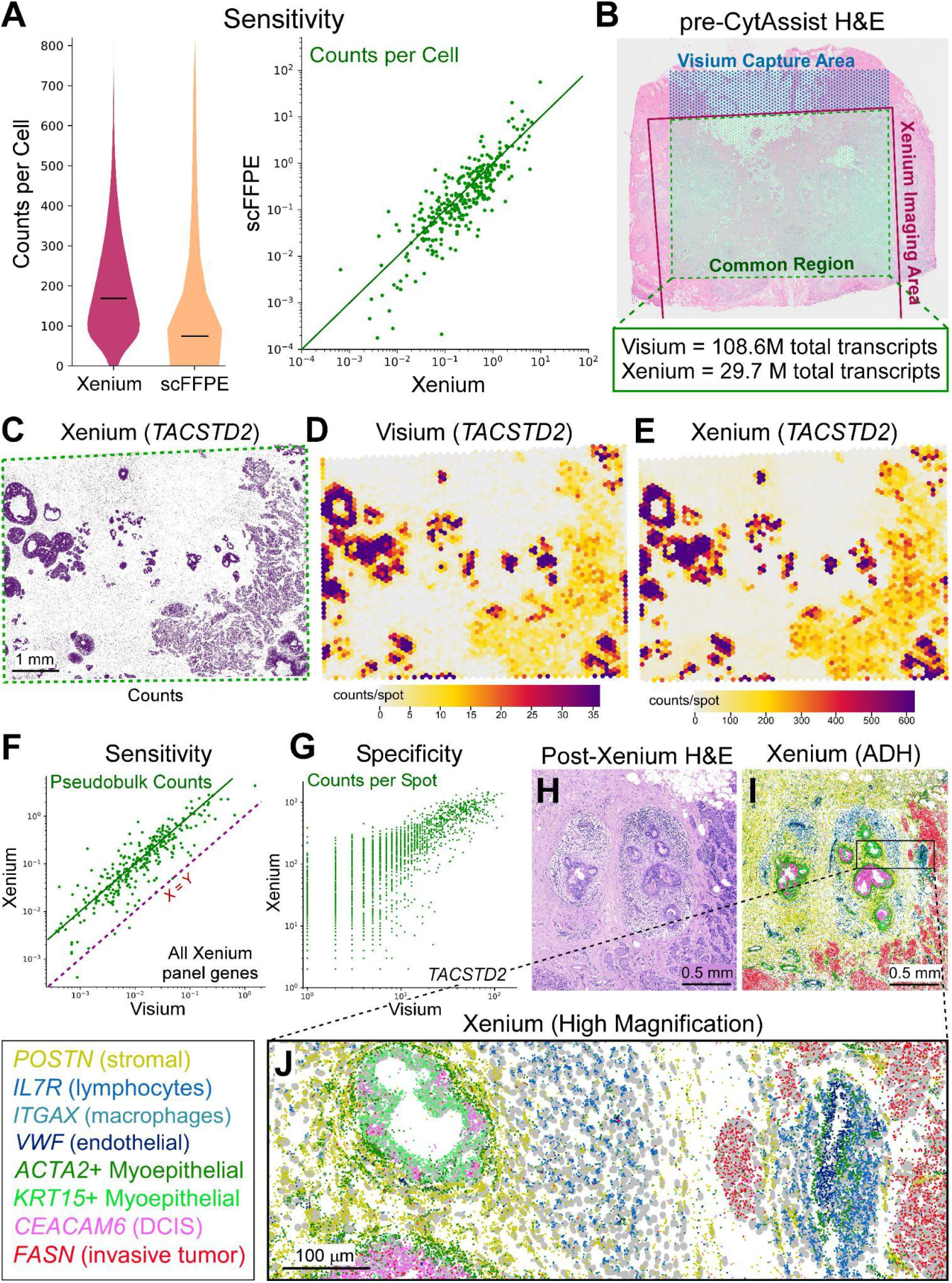
Xenium, Visium and scFFPE-seq are benchmarked for specificity, sensitivity, and resolution. (A) Chromium scFFPE-seq data down-selected for only the 313 genes that appear on the Xenium gene panel. Violin and scatter plots showing the total number of transcript counts per cell detected in the Xenium data vs. UMIs per cell in the scFFPE-seq data. (B) Registration of Visium and Xenium H&E allows for the isolation of a common area between the two platforms. The number of total transcripts within this common area is reported. (C-E) Tumor epithelial marker *TACSTD2* gene expression is shown as (C) Xenium transcript localization plot (decoded at Q ≥ 20), (D) Visium spatial plot (counts/spot), and (E) pseudobulk Xenium expression counts mapped to Visium spots. Resolution improvements from Visium are notable in the Xenium data, even at low magnification (compare C and D). (F) Sensitivity scatter plot expressed as pseudobulk counts per spot, quantified for all 313 genes on the Xenium panel. Dotted line represents X=Y; solid line is 10X = Y. (G) Comparison of Xenium and Visium data (counts per spot) showing high spatial correspondence (r^2^ = 0.88). (H) One small region of interest (ADH; atypical ductal hyperplasia) with a high diversity of cell types in close proximity is viewed with H&E staining. (I) Xenium spatial plot showing decoded transcripts for selected genes. Corresponding Visium spatial plots of the same genes are available in Supp. Fig. 9. (J) Closer view to reveal detailed resolution of Xenium nuclei (gray) and selected transcripts.

Next, we compared Visium and Xenium data by registering the corresponding H&E images to identify the common capture area (78% of the full Visium dataset) (Fig. 4B), which we focused on for subsequent analyses. Since Visium probes the whole transcriptome and Xenium probes 313 genes, Visium exhibited 3.6x more total transcripts within the shared region (Fig. 4B). Visium and Xenium exhibited concordant spatial expression (Supp. Fig. 8), exemplified by the tumor-associated epithelial marker *TACSTD2* (Fig. 4C, D). We mapped Xenium expression data onto the Visium capture area using the H&E registration information, and calculated pseudobulk counts within each Visium spot (Fig. 4E). The median gene sensitivity of Xenium across all genes on the human breast probe panel compared to Visium was 8.4x higher (Fig. 4F). To examine specificity, we compared *TACSTD2* transcript counts for Visium and Xenium and observed strong correlation (r^2^ = 0.88) (Fig. 4G), despite the increased sensitivity of the Xenium platform. While comparing individual genes allowed us to benchmark the Xenium workflow, the real power of in situ analysis is the ability to spatially localize multiple genes simultaneously at high resolution. This is especially useful in cases where many cell types coexist in close proximity such that they are not resolvable by other scRNA-seq and current capture-based spatial transcriptomics methods. When we examined a region of atypical ductal hyperplasia (ADH) in this tissue, we were able to clearly localize key markers for eight different cell types (Fig. 4H-J; Supp. Fig. 9).

### The Xenium workflow preserves epitopes, allowing RNA and protein to be visualized simultaneously in the same tissue section

Although there are some technologies available that analyze RNA and protein expression together, few methods allow for simultaneous visualization of both analytes on the same section at scale. Xenium on-instrument biochemistry and decoding cycles preserve protein epitopes, allowing for downstream IF (Fig. 5). The decrosslinking protocol for FFPE aids not only in RNA accessibility for the DNA probes, but also serves as an antigen retrieval step. Autofluorescence quenching (see Methods) substantially improves the signal-to-noise ratio. We demonstrated that protein epitopes are preserved by conducting standard immunofluorescence staining, after the Xenium workflow, on the same section. HER2 (tumor) and CD20 (B cell) antibodies were detected with fluorophore-conjugated secondary antibodies and their expression was compared to their cognate RNA (Fig. 5A, B). Their spatial expression was highly correlated, further highlighting the specificity of Xenium probes, optimized biochemistry, and decoding. Next, we registered the two maximum intensity projection DAPI images (Xenium on-instrument and immunofluorescence Zeiss microscope) to each other and overlaid protein and RNA data together (Fig. 5C-E). Because both images were taken from the same section, we were able to obtain a high degree of concordance and registration between the RNA and protein expression profiles.

**Figure 5.**
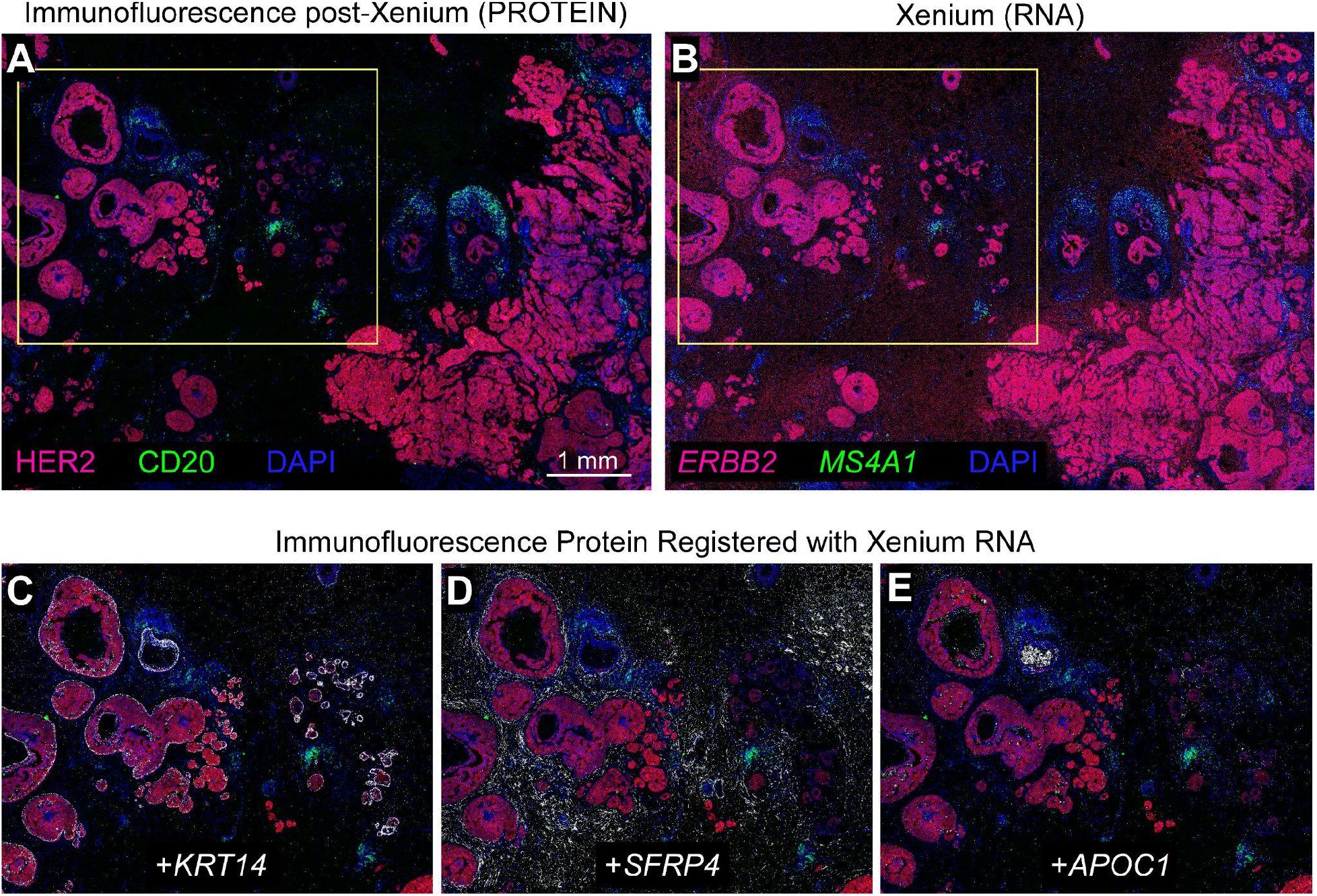
RNA from Xenium and protein immunofluorescence can be visualized in the same tissue section. (A) After 15 cycles of imaging on the Xenium instrument, sections were immunostained for HER2 and CD20 proteins using fluorescent secondary antibody detection in the 594 and 488 channels, respectively. (B) Comparing the equivalent Xenium RNAs (*ERBB2* and *MS4A1*), spatial correlation to protein expression was nearly identical. (C-E) Region of interest outlined with yellow box in (A) and (B). Because RNA and protein data were obtained from the same section, the two DAPI images were registered and overlaid with RNA and protein expression. HER2 and CD20 protein immunofluorescence are shown registered with (C) *KRT14* RNA (myoepithelial), (D) *SFRP4* RNA (stromal), and (E) *APOC1* RNA (macrophages).

### Chromium and Xenium integration explores the FFPE tumor microenvironment through cell type composition and differential gene expression analysis

Ductal carcinoma in situ (DCIS) is a non-obligate precursor of invasive ductal carcinoma, which can develop into invasive disease, the treatment of which often involves surgical removal of the lesion and radiotherapy (Wilson et al. 2022). Because not all DCIS lesions progress to invasive disease, there is great interest in understanding the molecular mechanisms underpinning invasiveness in DCIS, which are currently not well known (Wilson et al. 2022, Rebbeck et al. 2022), but could help to guide better therapeutic strategies. Our goal was to use Xenium and scFFPE-seq data to identify different tumor subtypes and supplement H&E imaging and pathology with molecular targets. First, we used scFFPE-seq data to map three different tumor epithelial cell subtypes and two myoepithelial subtypes to our Xenium data. We selected three regions of interest (ROIs): DCIS #1, DCIS #2, and invasive tumor (Fig. 6A). We verified these regions with expert pathologists who observed that 1) post-Xenium H&E exhibited high quality morphology comparable to standard H&E, and 2) ROIs were either morphologically distinct or were surrounded by a unique microenvironment. The pathologist annotation showed that DCIS #1 ROI had a smaller and round intermediate nuclear grade ductal hyperplasia of solid type with cells that showed mild to moderate variability in size, shape and placement, variable coarse chromatin, and variably prominent nuclei. DCIS #2 ROI had invasive carcinoma lesions scattered throughout the stromal connective tissue, surrounding a large aggregate of highly proliferative ductal carcinoma in situ with two central comedo necrotic formations.

**Figure 6.**
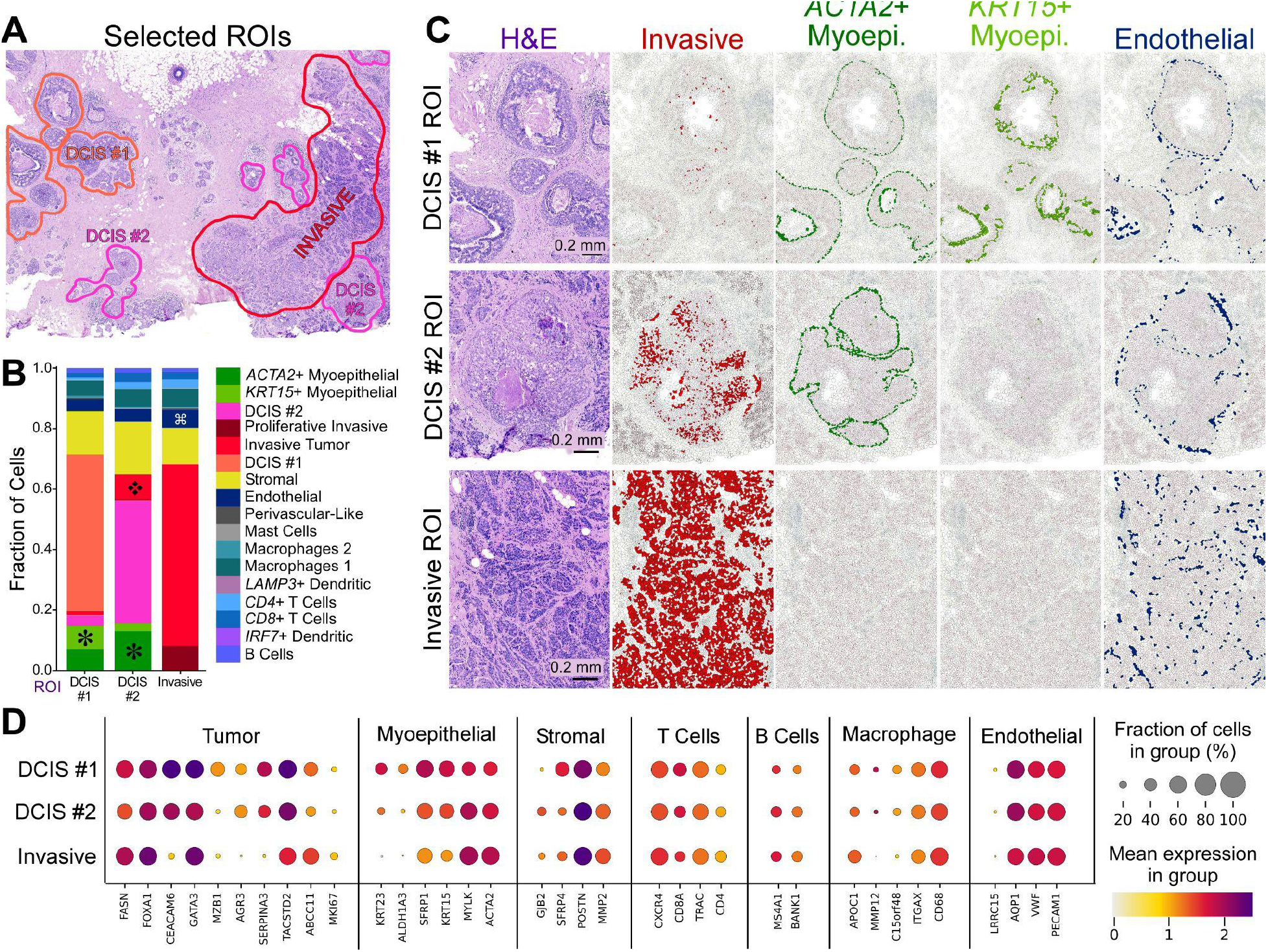
Integrating scFFPE-seq and Xenium data deciphers differences in cell type composition and molecular markers between DCIS subtypes and invasive tumor regions. (A) With pathology and scFFPE-seq guidance, we selected three ROIs capturing DCIS #1, DCIS #2, and invasive tumor cell types, and all other cell types in their proximity. (B) We determined the proportions of 17 cell types within these ROIs. We identified four major differences in cell type composition across the ROIs: ✽*ACTA2*+ and *KRT15*+ myoepithelial cell populations are distinct in DCIS #1 and DCIS #2 ROIs, but completely absent from invasive tumor ROI; ❖invasive tumor cells are found within the DCIS #2 ROI; ⌘ endothelial cells are found in slightly larger numbers within the invasive ROI. (C) Validation of the finding in (B). (D) Dot plots showing canonical markers of cell types as well as differentially expressed genes between the tumor subtypes.

We used scFFPE-seq data to determine proportions of 15 cell types within ROIs in the Xenium data, including lymphocytes, macrophages, stromal, myoepithelial, and invasive cells. We identified four major differences in cell type composition across ROIs (Fig. 6B). ACTA2+ myoepithelial cells were found to be prominent in DCIS #2 ROI, less common in DCIS #1 ROI, and absent in the invasive ROI, invasive tumor cells were found within DCIS #2 ROI, and endothelial cells were found in slightly larger numbers within the invasive ROI. We verified these findings (Fig. 6C) to illustrate how Xenium and scFFPE-seq data can uncover molecular differences which are not apparent with H&E pathology alone. The DCIS #2 ROI contained many more invasive cells than the DCIS #1 ROI, and also less *KRT15*+ myoepithelial cells, suggesting that DCIS #2 ROI is more invasive than DCIS #1 ROI. The invasive ROI had an extremely high incidence of invasive cell types, and the myoepithelial cell types were entirely absent. The high resolution of Xenium enables interactions among neighboring cells to be captured. This is well illustrated in the DCIS #2 ROI with the thin boundary of *ACTA2*+ myoepithelial cells encircling invasive cells (Fig. 6C).

Finally, we graphed the expression of canonical markers representing seven major cell types and differentially expressed genes between the tumor subtypes to provide insight into whether the DCIS ROIs were progressing to an invasive state (Fig. 6D). These analyses revealed that *MZB1* is an exclusive marker of the DCIS #1 ROI and cell type, *GJB2*+ stromal cells were found in the DCIS #2 ROI, *ALDH1A3*, *KRT15*, and *KRT23* were highly expressed in myoepithelial cells of the DCIS #1 ROI, and the macrophage marker *MMP12* was absent from the invasive ROI.

### Visium and Xenium integration derive whole transcriptome information from biological regions of interest

The hormone receptor status of a tumor is important biologically and has clinical relevance. The hormones estrogen and progesterone play a role in the development of breast epithelium during puberty, and exposure to ovarian steroids correlates with breast cancer risk (reviewed by Pal et al. 2021). Clinically, breast cancers are classified based on the expression of the estrogen receptor (ER / *ESR1*), progesterone receptor (PR / *PGR*), and human epidermal growth factor receptor 2 (HER2 / *ERBB2*) (Wu et al. 2021). These classifications typically define treatment strategies; for example, endocrine therapies are commonly used to treat patients with ER+ breast cancers (Coates et al. 2015). The tissue block used in this study was annotated as HER2+/ER+/PR−. The Xenium data shows mostly regions of *ERBB2+* (HER2+) and double positive *ERBB2+/ESR1+* (HER2+/ER+) gene expression (Fig. 7A). However, there is a small DCIS region located within an adipocyte region that is triple positive *ERBB2+/ESR1+/PGR+* (HER2+/ER+/PR+). Zooming in on this triple positive ROI with Xenium, H&E, and cell type labeling, we found a predominantly DCIS #2 tumor epithelium without a *KRT15*+ myoepithelial cell layer (Fig. 7B-D). Next, we compared expression of the three hormone receptor genes between the Xenium and scFFPE-seq data (Fig. 7E-F). While few *PGR*+ cells were found in the scFFPE-seq data, they did not seem to coincide with *ESR1* or *ERBB2* expression. In the Visium data, this region is represented by only 5-6 spots (Fig. 7G), which may have gone unnoticed. However, to derive whole transcriptome information from this triple positive region Visium proved critical, allowing us to identify the triple positive region as part of cluster 12, with five spots found in the adipocyte region (Fig. 7H). Using the spot interpolation methods (Supp. Fig. 10), we visualized the cell type proportions within this triple positive region, and selected the six spots that contain the DCIS #2 cell type (Fig. 7I). We then performed whole-transcriptome differential gene expression analysis of these six spots compared to all other Visium spots. This allowed us to identify 94 differentially expressed genes (log_2_FC >1.5; p-value < 0.05); four are shown here (Fig. 7J).

**Figure 7.**
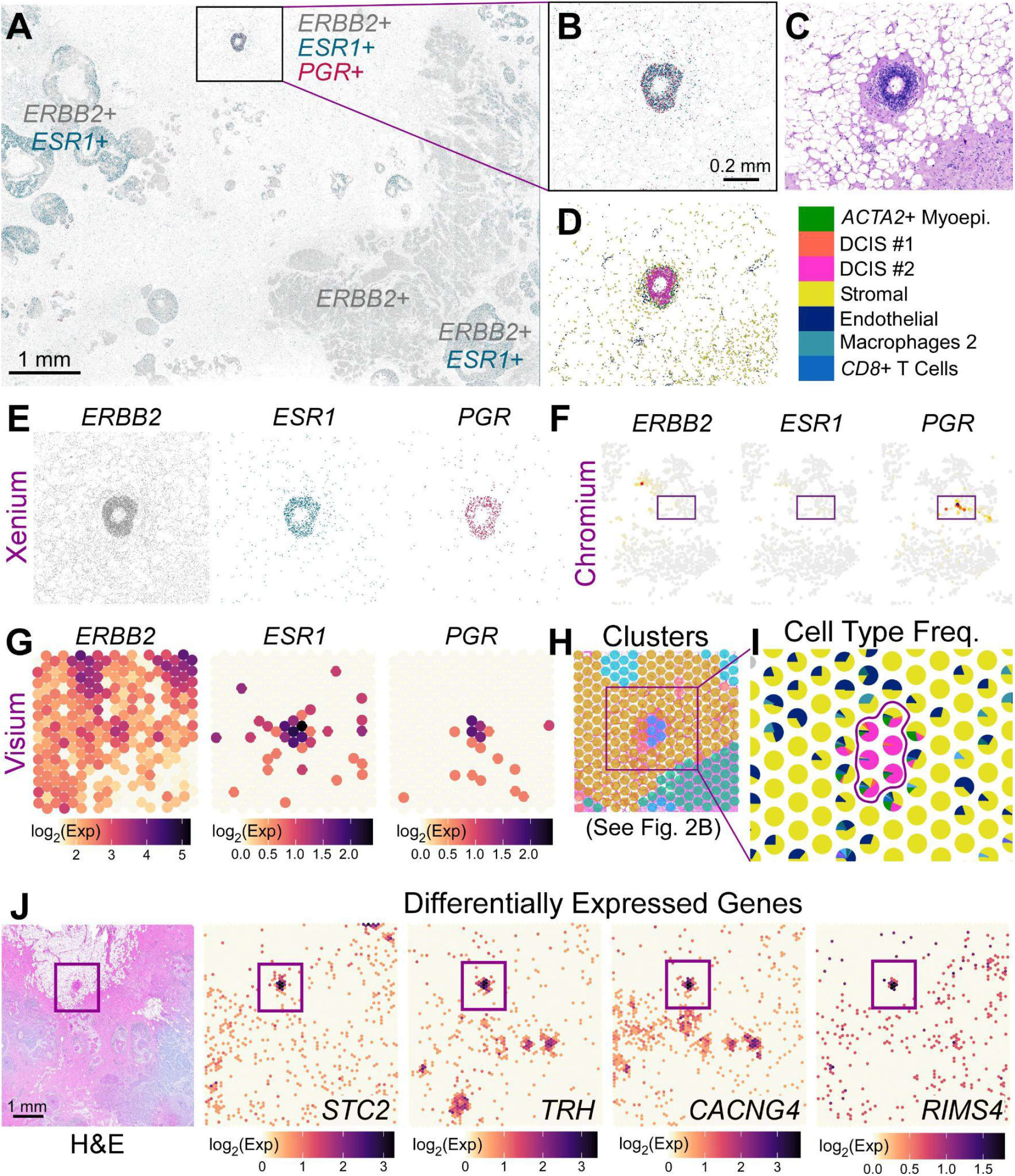
Visium and Xenium integration derive differentially expressed genes in a triple-positive receptor ROI. (A) Xenium spatial plot for *ERBB2* (HER2 - gray), *ESR1* (estrogen receptor - green), and *PGR* (progesterone receptor - magenta) decoded transcripts. (B) Closer view of triple-positive ROI. (C) Corresponding H&E image. (D) Cell types contained within ROI reveal that this is a DCIS #2 tumor epithelium. (E) Individual Xenium spatial plots from (B). (F) Chromium scFFPE-seq yields only about 30 cells that are positive for *PGR*, but these cells do not express *ERBB2* or *ESR1*. (G) Triple-positive region is identified in Visium (given *a priori* knowledge from Xenium) and is (H) part of a distinct cluster (see Fig. 2B). (I) Spot interpolation (see Supp. Fig. 10) provides cell type frequencies within each Visium spot. Color code legend is shown in (D). (J) Visium H&E and four representative differentially expressed genes in the tumor epithelium (94 genes; log_2_FC >1.5; p-value < 0.05) revealed by Visium data across the whole transcriptome.

## Discussion

Resolving the complexities of the tumor microenvironment is necessary for a comprehensive understanding of cancer biology. This is illustrated in our study using an FFPE block from a patient breast biopsy that contains both ductal carcinoma in situ and invasive ductal carcinoma. In this model, DCIS refers to neoplastic epithelial cells that remain confined within the ducts, and DCIS is therefore considered nonlethal (Allred 2010, Rebbeck et al. 2022, Wilson et al. 2022). DCIS can be (albeit is not always) the immediate precursor of potentially lethal invasive ductal carcinoma, when the ductal morphology is broken down and cancerous cells invade the stroma. Understanding why some DCIS regions become invasive, while others do not, remains an open question in the field. Attempts to answer this question often begin with clinical classifications of FFPE blocks by receptor type (e.g., HER2+/ER+/PR+) and degree of invasiveness and proliferation, but this taxonomy is insufficient to describe heterogeneity within the sample. Despite that the block used in our study was annotated by a pathologist as HER2+/ER+/PR−, we found a region of DCIS that was positive for the RNA of all three receptors, in just one 5.5 mm x 7.5 mm section of the much larger biopsy (Fig. 7). Furthermore, the FFPE block was annotated as 25% DCIS, but this did not capture the fact that at least two molecularly distinct DCIS regions were observed. One DCIS region showed an absence of myoepithelial markers along with the presence of cells already expressing an invasive molecular signature (Fig. 6), suggesting a progression to advanced invasive cancer.

Previous attempts to unmask such tumor heterogeneity, using bulk and single-cell next-generation sequencing (NGS) approaches and immunofluorescence, often target specific genes associated with an invasive/metastatic prognosis and treatment regime (Risom et al. 2022, Zhong et al. 2020, Garczyk et al. 2015). Our results are consistent with several published examples, albeit with higher resolution and gene plexy. For example, low keratin15 (*KRT15*) expression has been previously suggested to be associated with poor prognosis for patients with invasive carcinoma (Zhong et al. 2020). Our comparison of two DCIS regions with Xenium in situ data reveals reduced myoepithelial markers *KRT15*, *KRT23*, and *ALDH1A3* which could potentially be associated with increased invasiveness suggested by the higher expression of invasive markers found in DCIS #2 (Fig. 6). *ALDH1A3* (a.k.a. RALDH3), which catalyzes the formation of retinoic acid (RA), is pinpointed by Xenium to be highly expressed and spatially localized to the myoepithelial layer. Given the classical capacity for RA to behave as a morphogen, it would be important to understand where ALDH1A3 produces RA locally within the tissue. Although RA has varied roles in cancer, there is some evidence in cell lines that it increases senescence and adhesion to the basement membrane in breast myoepithelial cells, thereby decreasing the invasive capacity of tumor epithelium (Berardi et al. 2015). Xenium identified *AGR3* as a tumor epithelial marker associated with DCIS ROIs, but not the invasive ROI (Fig. 6). Likewise, Garczyk et al. (2015) found that *AGR3* protein expression in breast tumors is significantly associated with estrogen receptor α and lower tumor grade, suggesting that *AGR3* could serve as a biomarker for prognosis and early detection. This ability of Xenium to map the localization and expression level of key genes at high resolution holds great promise to transform, diagnose, prognose and guide more effective treatment management.

Female breast cancer is the most commonly diagnosed cancer globally, with ~2.3 million new cases reported in 2020 (Sung et al. 2021), and the prevalence of cancer as a leading cause of premature death is ever-increasing worldwide, particularly in developing nations (Bray et al. 2021). How can single-cell, spatial, and in situ technologies scale to deal with this challenge? The key to this question is the ability of these technologies to glean high quality data from FFPE tissues. FFPE methods for preservation of samples are well established in clinical practice as they allow for a high degree of morphological detail to be maintained, and as such, there are large numbers of FFPE specimens in biobanks that are potentially available for genomics research (Villacampa et al. 2021). Using carefully designed probe sets and RTL technology, we are able to overcome the formalin-induced obstacles of strand cleavage and cross-linking that have plagued researchers for decades.

In this study, we used these three independent but complementary genomics technologies to explore the biology of a single FFPE-preserved tissue block. Our results typify how the integration of these technologies is an iterative process, and suggest how discoveries in one data modality can rapidly inspire explorations in another. What did each technology bring to the table, and what did we learn from integrating them that we could not have learned from a single technology individually? scFFPE-seq is the most sensitive of the three, particularly for lowly expressed genes. We found that scFFPE-seq median gene sensitivity was higher than both Chromium 5′ and 3′ GEX data, from patient-matched dissociated tumor cells (Supp. Fig. 7). Of the three FFPE-compatible assays presented here, scFFPE-seq is the only one offering whole transcriptome data at single cell resolution, making it well suited for establishing a baseline of disease, annotating cell types (Fig. 2A), and designing or validating targeted Xenium gene panels. Like scFFPE-seq, Visium also provides whole transcriptome data. Although Visium lacks true single cell resolution at this time, it provides a spatial context that cannot be explored with single cell technologies. Integrating scFFPE-seq and Visium data was straightforward due to the identical probe set used in both technologies, and allowed for accurate deconvolution of cell types that composed the Visium spots (Supp. Fig. 10).

In the early stages of data exploration, Visium and H&E data were used to annotate three tumor cell types within the scFFPE-seq data by noting that the differentially expressed genes in specific scFFPE-seq clusters were mapping to the invasive tumor domain, or one of two spatially distinct DCIS regions in the H&E image (Fig. 2). Hence, Visium alone identified that there were three spatially distinct tumor subtypes, which was not captured in the pathologist annotations. We then integrated Xenium and Visium to derive differentially expressed genes from a tumor region containing cells expressing RNA of three receptors (ERBB22+/ESR1+/PGR+). Neither Visium nor scFFPE-seq identified these cells initially because they were so sparse, and required the high resolution spatial information gained by Xenium. Using spot interpolation methods (Supp. Fig. 10), we identified the relevant tumor epithelial cells in the Visium data and derived whole transcriptome information. Visium and Xenium data were also able to recover adipocytes (Fig. 2, Supp. Fig. 4), which typically are lost during the sample preparation protocols necessary for single cell analysis. Xenium is particularly suited for investigations of intricate tissues with a high diversity of cell types in a small area, including immune and myoepithelial cells (Fig. 4, Supp. Fig. 9), that may elude the other two technologies. Another advantage is that because the Xenium workflow has a relatively low impact to the tissue, H&E and IF can be performed after the gene expression data are collected (Figs. 3C, 4H, Fig. 5). The importance of having pathology information associated with the molecular information from the same section cannot be overstated. Even 5-10 μm serial sections can change drastically enough that, at a single cell level, the hematoxylin-stained and DAPI-stained nuclei cannot be easily overlain. H&E staining is well established among pathologists due to excellent contrast, color, and texture between various cell features, so having both H&E and molecular information is valuable. Similarly, the incorporation of an additional protein molecular layer on the same section as the RNA readout provides richer phenotypic data.

High resolution in situ analysis of complex tissues will revolutionize how we understand biology, providing insights not previously possible with other technologies. As the Xenium platform capabilities are built out further, they will include increased gene plexy, more tissue-specific gene panels, decoding of RNA and protein from the same tissue section, detection of single nucleotide polymorphisms and isoforms (Lebrigand et al. 2022), and continually improving analysis and software tools. This will lead to an even greater understanding of molecular profiles as they relate to the tissue architecture, and how cells interact with other cells and non-cellular components in their local tissue environment. Our findings here demonstrate that the highest resolution and richest biological information are gleaned through the combination of complementary technologies. While each technology independently elucidates high quality gene expression data from FFPE tissues, it is the integration that illuminates biology with more rigor and refinement than a single technology alone. The resolution and breadth of the platforms we describe have promising implications across the biological sciences, but particularly in the future of translational and clinical research, and ultimately, in advancing human health.

## Data Sharing / Accessibility

### Downloadable datasets

https://www.10xgenomics.com/products/xenium-in-situ/preview-dataset-human-breast

### Interactive data explorer

https://www.10xgenomics.com/products/xenium-in-situ/human-breast-dataset-explorer

## Methods

### Samples and sample collection

A single formalin-fixed, paraffin-embedded (FFPE) breast cancer tissue block (TNM stage T2N1M0, ER+/HER2+/PR−) was collected on 2021-07-26 and obtained from Discovery Life Sciences. Corresponding dissociated tumor cells, fresh frozen in liquid nitrogen, were also sampled from the same biopsy (patient matched). 5 μm sections were taken from the FFPE tissue using a microtome (Thermo Scientific HM355S; MX35 blades). For the Chromium Fixed RNA Profiling (scFFPE-seq) workflow, 25 μm FFPE curls were collected into a tube prior to serial sectioning for Visium CytAssist and Xenium (two replicates of 5 μm sections for each spatial platform), then an additional 25 μm FFPE curl was collected into the same tube reserved for scFFPE-seq. These pooled 25 μm curls (50 μm total) were treated as a single replicate.

### Chromium 3′ and 5′ Single Cell Gene Expression (GEX)

We collected Chromium 3′ and 5′ GEX data from dissociated tumor cells to benchmark performance against the scFFPE-seq data. Dissociated tumor cells were recovered following Demonstrated Protocol CG000233. For the 3′ and 5′ workflows, cells were loaded on to the Chromium X instrument following the library preparation protocols in the Chromium Next GEM Single Cell 3’ Reagent Kits v3.1 User Guide (CG000204) and Chromium Next GEM Single Cell 5’ Reagent Kits v2 (Dual Index) User Guide (CG000331), respectively. Libraries were sequenced on an Illumina NovaSeq with paired-end dual-indexing (28 cycles Read 1, 10 cycles i7, 10 cycles i5, 90 cycles Read 2). All of the 3′ and 5′ flowcells were demultiplexed with bcl2fastq (Illumina). FASTQ files were processed with Cell Ranger v7.0.1 (10x Genomics), using the cellranger count pipeline on each GEM well with the GRCh38-2020-A reference to produce gene-barcode matrices and other output files, followed by aggregation of GEM wells with the cellranger aggr pipeline.

### Chromium Fixed RNA Profiling (scFFPE-seq)

Our goal in producing scFFPE-seq data was to precisely define the cell types present in serial tissue sections to enable downstream integration of data types. 50 μm FFPE curls were dissociated with the Miltenyi Biotech FFPE Tissue Dissociation Kit. Approximately 600,000 cells were washed, counted and resuspended, loading 16,000 cells per each of four GEM wells (targeting 10,000 recovered cells) on a single Chromium X chip. Sequencing libraries were generated following the Chromium Fixed RNA Profiling for Singleplexed Samples User Guide (CG000477). Libraries were sequenced on an Illumina NovaSeq with paired-end dual-indexing (28 cycles Read 1, 10 cycles i7, 10 cycles i5, 90 cycles Read 2). Sequencing libraries were demultiplexed with bcl2fastq (Illumina). FASTQ files were processed with Cell Ranger v7.0.1 (10x Genomics) using the multi pipeline and the GRCh38-2020-A reference.

### Visium CytAssist

Whole transcriptome spatial data. Our goal in producing Visium CytAssist data was to obtain whole transcriptome, spatially-barcoded sequence data in serial sections. The histology workflow was performed using the Visium CytAssist Spatial Gene Expression for FFPE (Demonstrated Protocol CG000520). The tissue was sectioned as described in Visium CytAssist Spatial Gene Expression for FFPE – Tissue Preparation Guide (Demonstrated Protocol CG000518). 5 μm sections were placed on a Superfrost™ Plus Microscope Slide (Fisherbrand™) and H&E-stained following deparaffinization. Sections were imaged, decoverslipped, followed by hematoxylin destaining and decrosslinking (Demonstrated Protocol CG000520). The glass slide with tissue section was processed with a Visium CytAssist instrument to transfer analytes to a Visium CytAssist Spatial Gene Expression slide with a 0.42 cm^2^ capture area. The probe extension and library construction steps follow the standard Visium for FFPE workflow outside of the instrument. Libraries were sequenced with paired-end dual-indexing (28 cycles Read 1, 10 cycles i7, 10 cycles i5, 90 cycles Read 2). Sequencing libraries were demultiplexed with bcl2fastq (Illumina). The Space Ranger pipeline v2022.0705.1 (10x Genomics) and the GRCh38-2020-A reference were used to process FASTQ files.

### Xenium In Situ Workflow

#### Gene Panel Design

The Xenium In Situ technology uses targeted panels to detect gene expression. 313 genes for cell type identification (280 of which are included in the Xenium Human Breast Panel) were selected and curated primarily based on single cell atlas data for human breast tissue (Pal et al. 2021, Bhat-Nakshatri et al. 2021, Karlsson et al. 2021). The probes were designed to contain two complementary sequences that hybridize to the target RNA and a third region encoding a gene-specific barcode, so that the paired ends of the probe bind to the target RNA and ligate to generate a circular DNA probe. If the probe experiences an off-target binding event, ligation should not occur, suppressing off-target signals and ensuring high specificity.

#### Xenium Sample Preparation

The Xenium workflow (using in-development chemistry and a prototype instrument) began by sectioning 5 μm FFPE tissue sections onto a Xenium slide, followed by deparaffinization and permeabilization to make the mRNA accessible. The mRNAs were targeted by the 313 probes described above and two negative controls: 1) probe controls to assess non-specific binding and 2) genomic DNA (gDNA) controls to ensure the signal is from RNA. Probe hybridization occurred at 50° C overnight with a probe concentration of 10 nM. After stringency washing to remove un-hybridized probes, probes were ligated at 37° C for two hours. During this step, a rolling circle amplification (RCA) primer was also annealed. The circularized probes were then enzymatically amplified (for one hour at 4° C followed by two hours at 37° C), generating multiple copies of the gene-specific barcode for each RNA binding event, resulting in a strong signal-to-noise ratio. After washing, background fluorescence was quenched chemically. The biochemistry is designed to mitigate autofluorescence, which is a known issue due to the presence of lipofuscins, elastin, collagen, red blood cells, and formalin-fixation itself (Davis et al. 2014). Sections were placed into an imaging cassette to be loaded onto the Xenium Analyzer instrument.

#### Xenium Analyzer Instrument

The Xenium Analyzer is fully automated and includes an imager (imageable area of about 12 x 24 mm per slide), sample handling, liquid handling, wide-field epifluorescence imaging, capacity for two slides per run, and an on-instrument analysis pipeline. The imager is a fast area scan camera featuring a high numerical aperture, a low read noise sensor, and ~200 nm per-pixel resolution. On the Xenium Analyzer, image acquisition was performed in cycles. The reagents, including fluorescently labeled probes for detecting RNA, were automatically cycled in, incubated, imaged, and removed by the instrument. Following the binding of fluorescent oligos to the amplified barcode sequence, the sample underwent 15 rounds of fluorescent probe hybridization, imaging, and probe removal. The Z-stacks were taken with a 0.75 μm step size across the entire tissue thickness.

#### Image pre-processing

The Xenium Analyzer captured a Z-stack of images every cycle and in every channel, which needed to be processed and stitched to build a spatial map of the transcripts across the tissue section. Stitching was performed on the DAPI image, taking all of the stacks from different FOVs and colors to create a single seamless image representative of one ROI.

#### Puncta detection

A punctum (plural: puncta) is a point source in microscopy, smaller than a pixel, and is measured in units of observed photons. The pipeline detected every punctum, in every cycle, every image, every color, in order to observe all potential mRNA. First, puncta were localized by fitting a Gaussian distribution to the observed emitted light to determine the center, size, and intensity of the point sources. Next, the puncta in different cycles were registered to one another so they are occupying the same space (the same mRNA molecule emitting light).

#### Decoding and quality scores

In each cycle, fluorescently-labeled oligonucleotides are bound to amplified barcodes and the fluorescent intensity in each of the four Xenium color channels is measured. Using the fluorescent intensity detected over the 15 Xenium cycles, an optical signature unique for each gene is generated and used to identify a target gene. Each decoded transcript was assigned a raw Q-score, a phred-scaled Quality Value which indicates the probability that the detected object exists and was correctly identified by the decoding algorithm. The Q-score is influenced by many technical factors such as signal brightness, spot localization accuracy, and signal purity. Negative control codewords built into the system ensure that the reported Q-score is accurately calibrated. All Xenium spatial gene plots shown are transcripts passing Q ≥ 20.

#### Cell segmentation

In order to assign mRNAs to cells, and thus enable downstream analysis and integration with Chromium and Visium data, the spatial boundaries of cells relative to mRNA transcripts must be defined. First, DAPI images were used to detect nuclei using a neural network. Then each nucleus was expanded outwards until either 15 um max distance was reached or the boundary of another cell was reached.

#### Output file export

A variety of output files were produced by the on-instrument pipeline. The essential files used downstream were the feature-cell matrix (HDF5 and MEX formats identical to those output by Cell Ranger and Space Ranger for Chromium and Visium data, respectively), the transcripts (listing each mRNA, its 3D coordinates, and a quality score), and the cell boundaries CSV file. These files were then transferred for downstream analysis off-instrument.

### Post-Xenium Histology

#### H&E and IF staining

The post-Xenium H&E staining followed Demonstrated Protocol CG000160. For post-Xenium IF staining, sections were washed with PBST, then incubated in a blocking buffer (ScyTek AAA999) for 30 minutes at room temperature. The primary antibody in the blocking buffer was added and incubated in the dark at 4 C overnight. The following day, the sections were washed three times (10 minutes each) with PBST then incubated with secondary antibodies and DAPI in a blocking buffer, in the dark at room temperature, for two hours. Next, the sections were washed three times (10 minutes each) with PBST. Sections were imaged in a proprietary Xenium imaging buffer and imaged on a Zeiss Axioimager with a 40x dipping objective. The Zen Blue software was used for tiling, image acquisition, and exporting TIFF files. Post-Xenium H&E and IF images were registered to Xenium data using a custom script.

| Protein | Host | Fluor | Vendor | Catalog | Dilution |
| --- | --- | --- | --- | --- | --- |
| CD20 | Mouse | Unconj. | Abcam | AB219329 | 1:200 |
| HER2 | Rabbit | Unconj. | Abcam | AB134182 | 1:1500 |
| Ms IgG | Goat | Alexa 488 | ThermoFisher | A-11029 | 1:500 |
| Rb IgG | Goat | Alexa 594 | Abcam | AB150088 | 1:500 |

### Downstream analysis & integration

#### Chromium & Visium post-processing

The 3′, 5′, and scFFPE-seq data were filtered with scanpy 1.19. Cell filtering parameters included largest gene fraction ≤ 0.2, mitochondrial fraction ≤ 0.15, and number of genes observed ≥ 500. We performed t-distributed stochastic neighbor embedding (t-SNE) on Chromium and Xenium data using the monet package (Wagner 2020). A principal component analysis (PCA) was performed on the feature-cell matrix, and the top 50 components were input to the t-SNE. We subsampled the whole transcriptome Chromium and Visium data to only the 313 genes used in the Xenium panel. The Xenium t-SNE coordinates were initialized with the scFFPE-seq cluster centers. The Visium t-SNE was generated in Loupe, and expression is either reported as log_2_(counts) when shown stand-alone, or as raw counts when comparing directly with Xenium transcript counts.

#### Supervised labeling & label transfer

We annotated the scFFPE-seq data by first conducting a differential gene expression (DGE) analysis across unsupervised clusters in Loupe. Annotations were built upon this DGE analysis and literature review (Wu et al. 2021, Karlsson et al. 2021). We performed a log-normalization step of the data, and then calculated a z-score across cells. A PCA was performed and the top 50 PCs were selected. From the in situ data, we determined the 30 nearest neighbors for each cell after normalization and projection into PC space, and if at least 50% were one cell type, then that is the cell type that was assigned. If that criteria was not met, then the cell was classified as “unlabeled”.

#### Xenium Differential Gene Expression (DGE)

We drew a region of interest (ROI), a polygon around morphological features (individual cells, groups of cells, etc.) and performed DGE across these ROIs with scanpy v1.19. ROI selection was performed in the Xenium Explorer software (development version, 10x Genomics), and significance was assessed with the Wilcoxon test on log-normalized count data. The DGE was performed for each cell type across ROIs.

#### Benchmarking sensitivity

Because mean sensitivity is biased by high expressors, we calculated median gene sensitivity by first computing the sensitivity of each gene separately (the mean of the counts per cell), then calculating the median across all genes. Because sensitivity is dependent on sequencing saturation, the 3′ and 5′ GEX data were downsampled to 10,000 mean reads per cell to match the sequencing depth of 10,000 reads per cell (the recommended depth) for scFFPE-seq, and 20,000 mean reads per cell (the recommended sequencing depth for the 3’ and 5’ assays). The 3′ and 5′ GEX data were also downsampled to only the genes on the RTL scFFPE-seq probe set.

#### Image registration

For registration of IF images to the Xenium morphology images, which are both DAPI images, we used a SIFT registration with the cv2 4.5.4 package in python v3.9.7, which produces the transformation between IF and Xenium. For registration of Visium to Xenium data, serial sections were rotated 2.58 degrees relative to each other, then a manual-defined keypoint registration between the corresponding H&E images (serial sections) was used. Over 100 landmark features were identified on commonly shared microstructures. Using RANSAC, we determined the subset of coordinates that matched, and performed the transformation between coordinates with the FindHomography() function in the cv2 package.

#### Visium / Xenium Spot interpolation and deconvolution

Using the registration of Xenium to Visium, we binned cells (by centroid) and transcripts from Xenium into the Visium spots. This was done by proximity. The closest spot to a cell or transcript was identified as the spot a cell or transcript lies within. Robust Cell Type Decomposition (RCTD) with spacexr 2.0.1 (Cable et al. 2021) in R was used to deconvolve Visium spots into cell types using the unsupervised scFFPE-seq reference.

## Supplemental Material

### Supplemental Figures

**Supplemental Figure 1.**
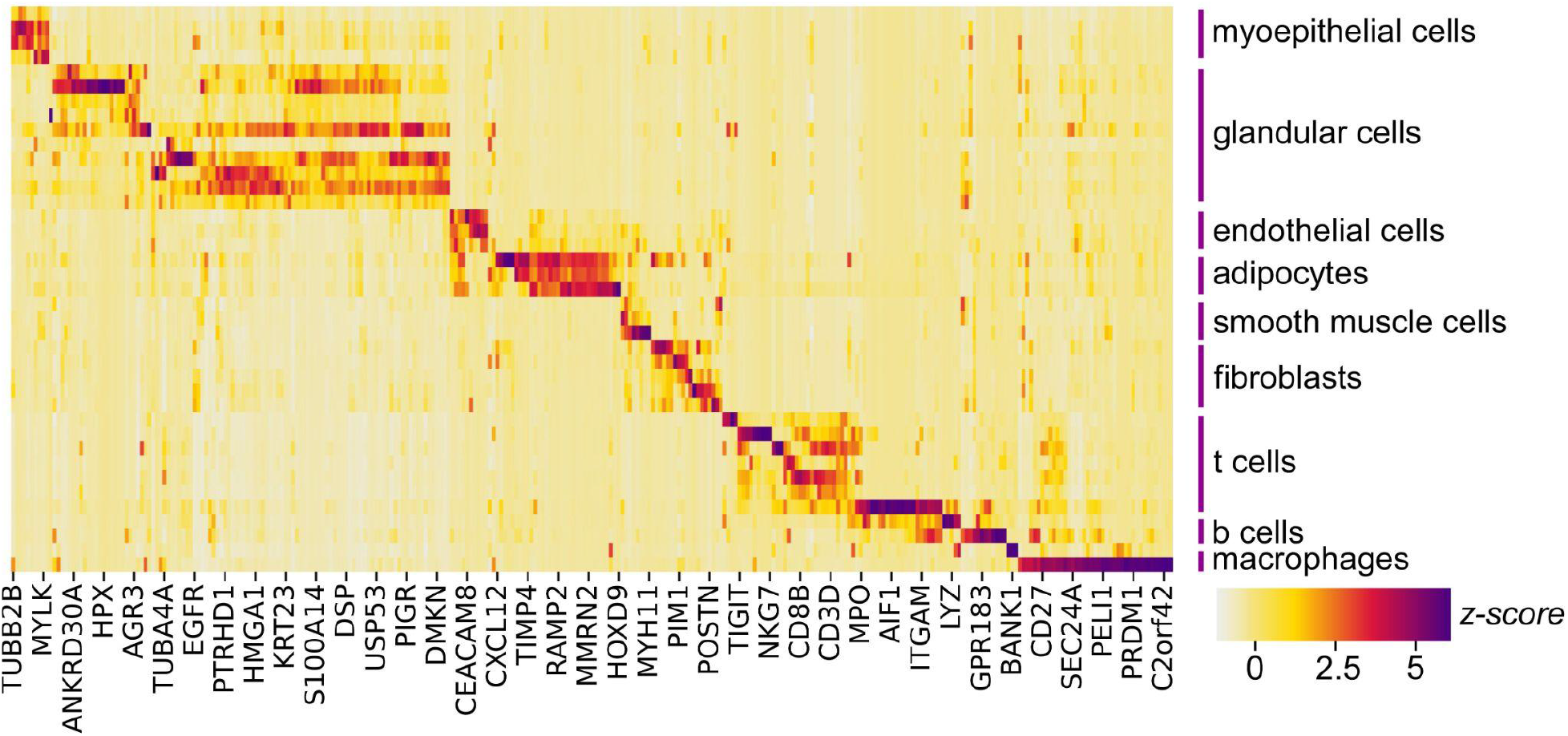
Xenium Gene Panel Design. The Xenium In Situ technology uses targeted panels to detect gene expression. The 280 gene pre-designed Xenium human breast panel was combined with 33 add-on genes, selected based on single cell data of human breast tissue (Pal et al. 2021, Bhat-Nakshatri et al. 2021, Karlsson et al. 2021) and manual curation. The heatmap shows the relative expression of genes across different cell types found in the references used to build the panel. The gene markers chosen are generally mutually exclusive for their cell type. Z-score computed across cell types for each gene.

**Supplemental Figure 2.**
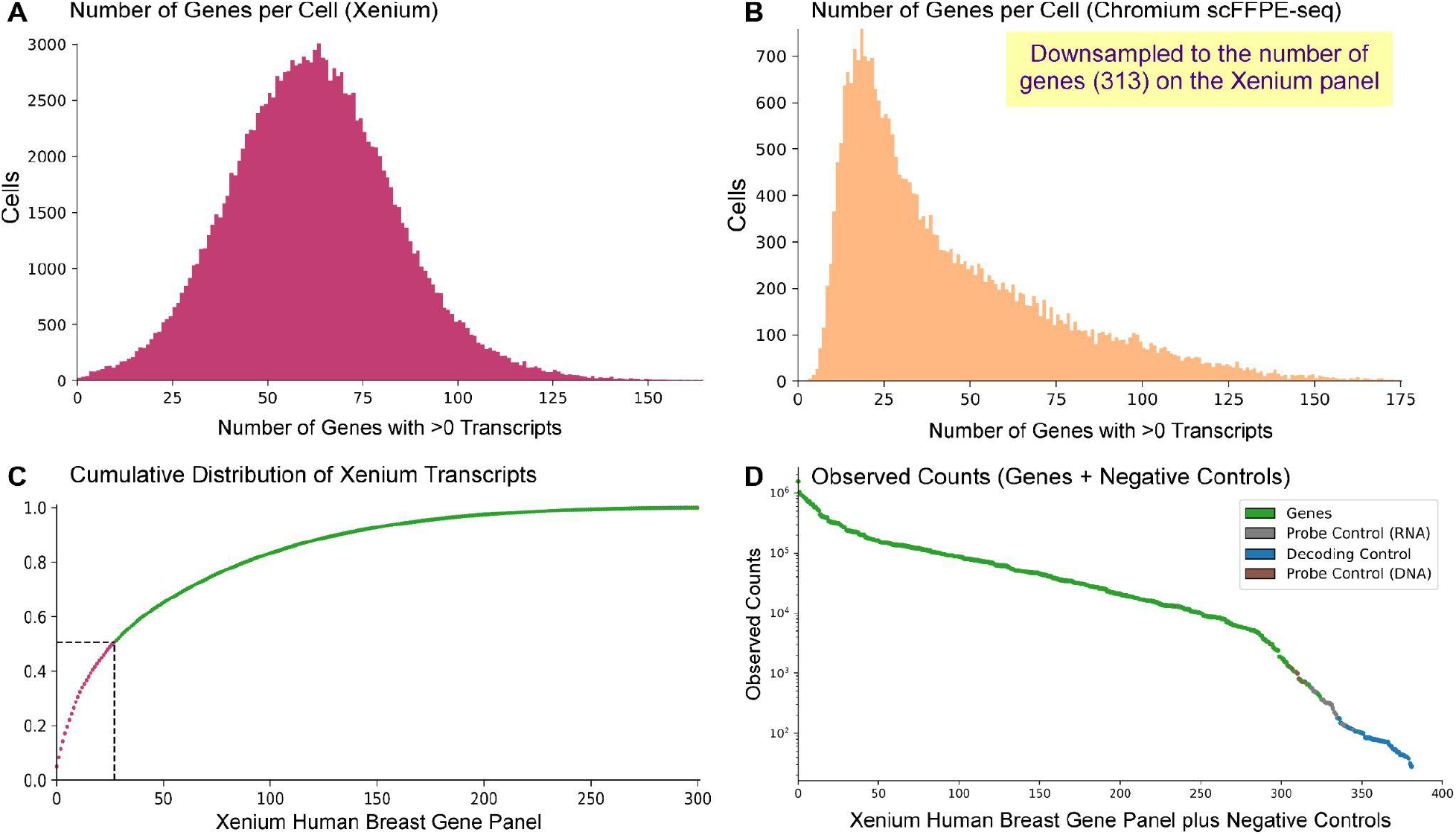
Xenium Quality Control Metrics. (A, B) Bar plots showing the number of genes detected per cell for (A) Xenium compared to (B) scFFPE-seq (downsampled to the 313 genes on the Xenium panel). See Fig. 3E for transcripts per cell. (C) Complexity measurement showing the cumulative distribution plot of total transcripts contributing to genes on the Xenium Human Breast Panel. Dotted line at y = 0.5 signifies that 50% of the total transcripts observed contribute to 27 genes. (D) Knee plot showing observed counts (Q ≥ 20) of genes and negative controls: 1) probe controls to assess non-specific binding to RNA, 2) decoding controls to assess misassigned genes, and 3) genomic DNA (gDNA) controls to ensure the signal is from RNA.

**Supplemental Figure 3.**
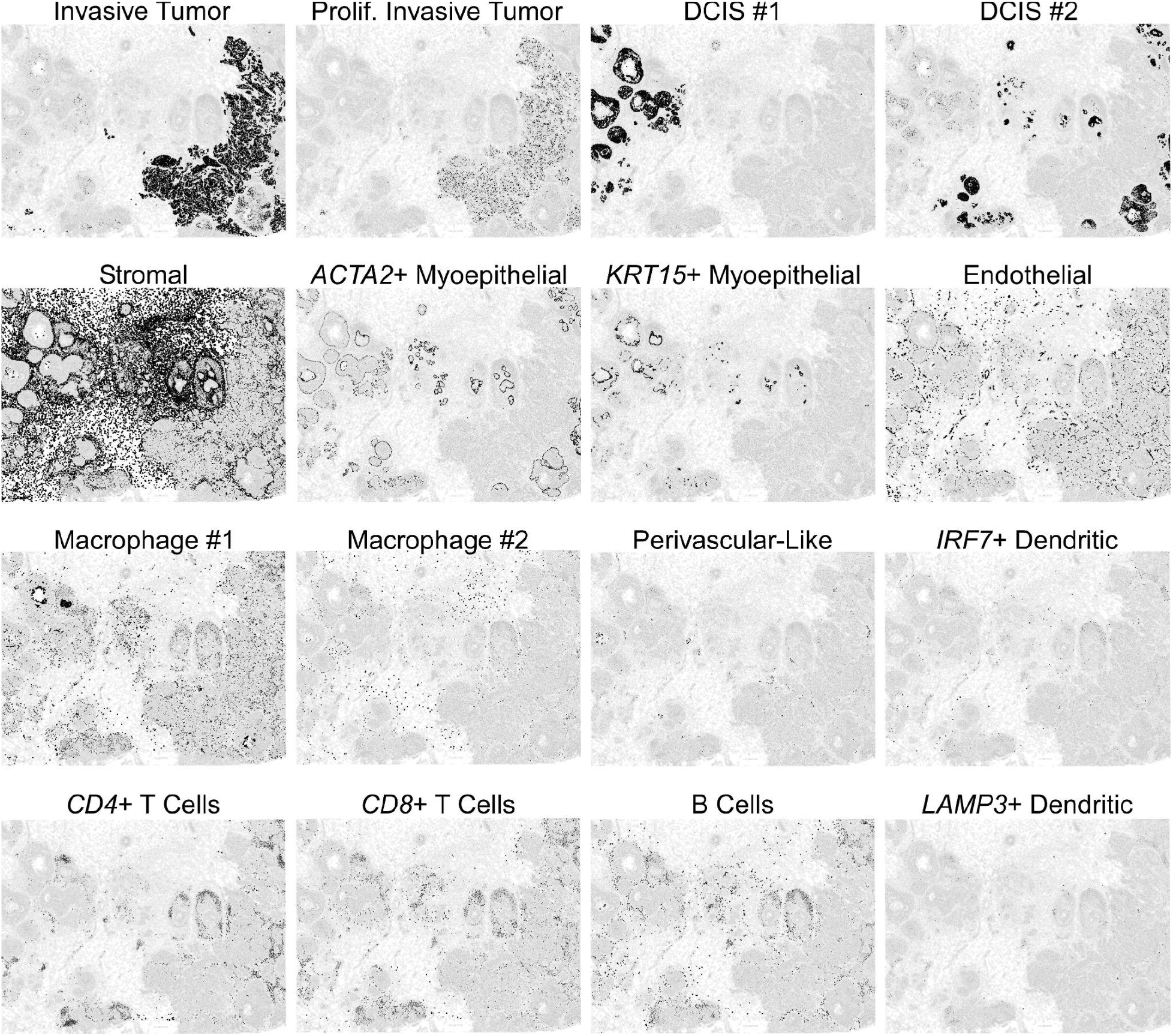
All cell types pictured in Fig. 3J are shown here individually. The highlighted cell type is filled black, and all other cells are filled white.

**Supplemental Figure 4.**
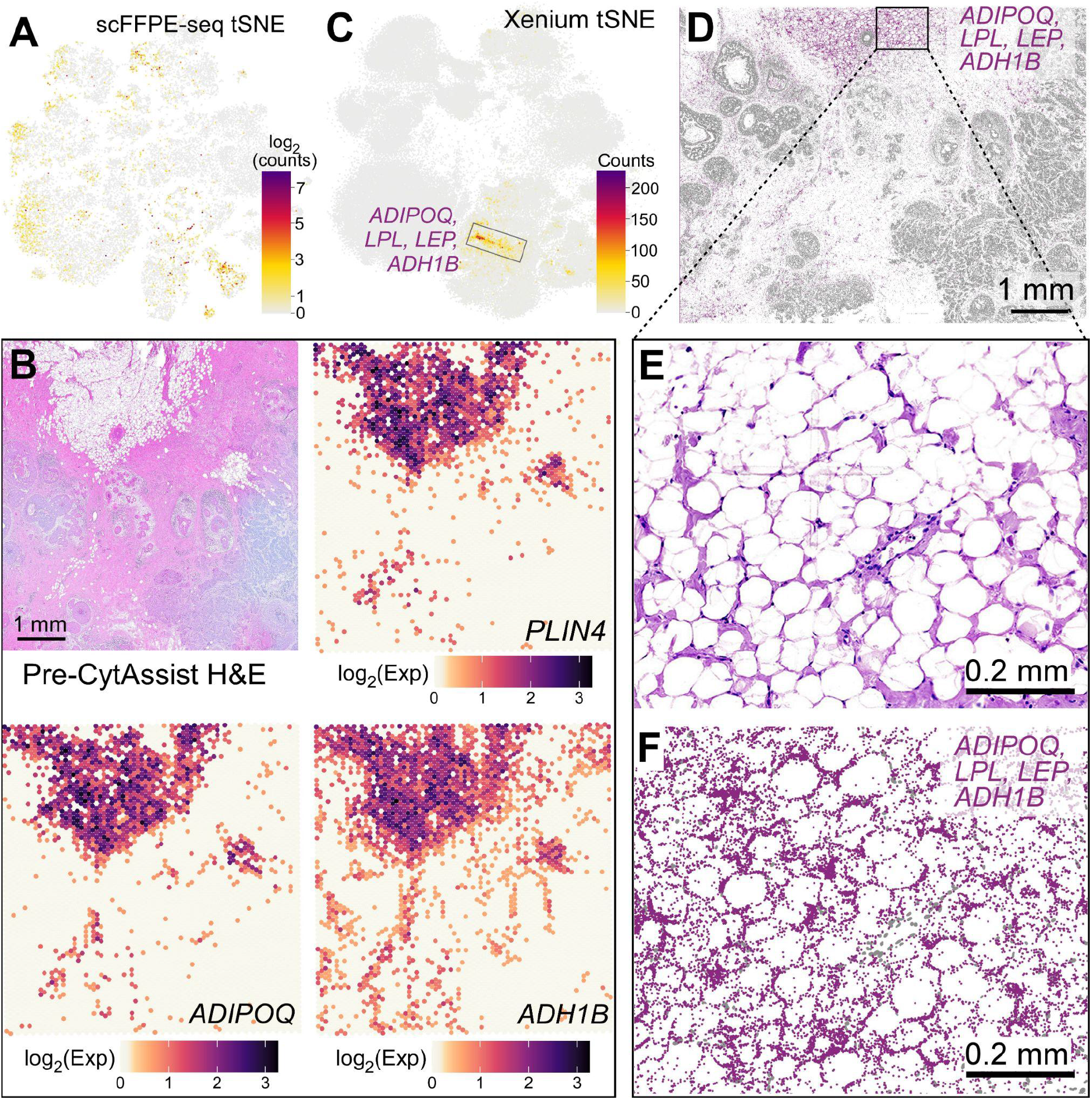
Visium and Xenium data identify adipocytes and their gene markers. (A) t-SNE projection of the scFFPE-seq data showing the combined expression of 11 adipocyte markers (*FABP4*, *GPX3*, *PLIN1*, *PLIN4*, *ADH1B*, *ADIPOQ*, *G0S2*, *LPL*, *GPD1*, *LIPE*, *SLC19A3*). See Fig. 2A for cluster annotations. (B) H&E staining conducted pre-CytAssist is shown for reference alongside the Visium spatial distribution of three known adipocyte markers (*PLIN4*, *ADIPOQ*, and *ADH1B*) expressed as log_2_(normalized UMI counts). (C) t-SNE projection and (D) spatial plot of the Xenium data displaying the expression of all adipocyte markers on the Human Breast Panel: *ADIPOQ*, *LPL*, *LEP*, and *ADH1B*. (E, F) Closer view of an adipocyte region showing (E) post-Xenium H&E and (F) Xenium spatial plot for adipocyte markers with nuclei in gray.

**Supplemental Figure 5.**
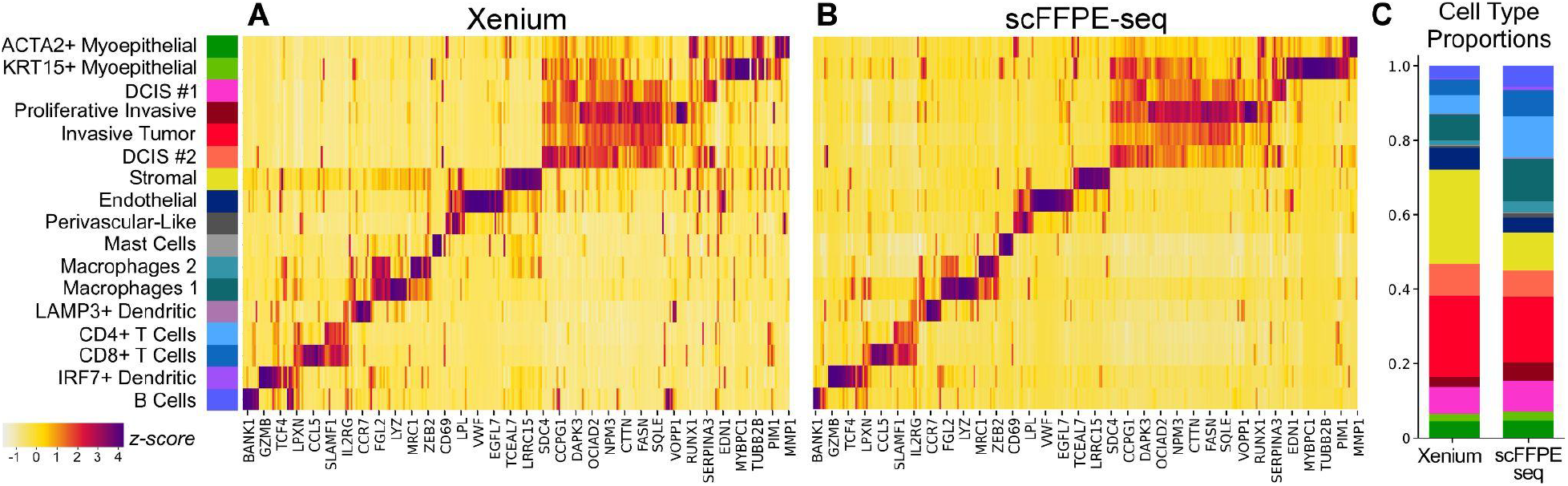
Relative expression of genes across cell types comparing Xenium and. scFFPE-seq. (A, B) Heatmap representation of the t-SNEs in Fig. 3H (Xenium) and Fig. 3G (scFFPE-seq, down-selected to 313 genes), demonstrating that the cell type representation is generally similar. Z-score computed across cell types for each gene, by subtracting the mean and dividing by the standard deviation. (C) Comparison of cell type proportions in Xenium versus scFFPE-seq.

**Supplemental Figure 6.**
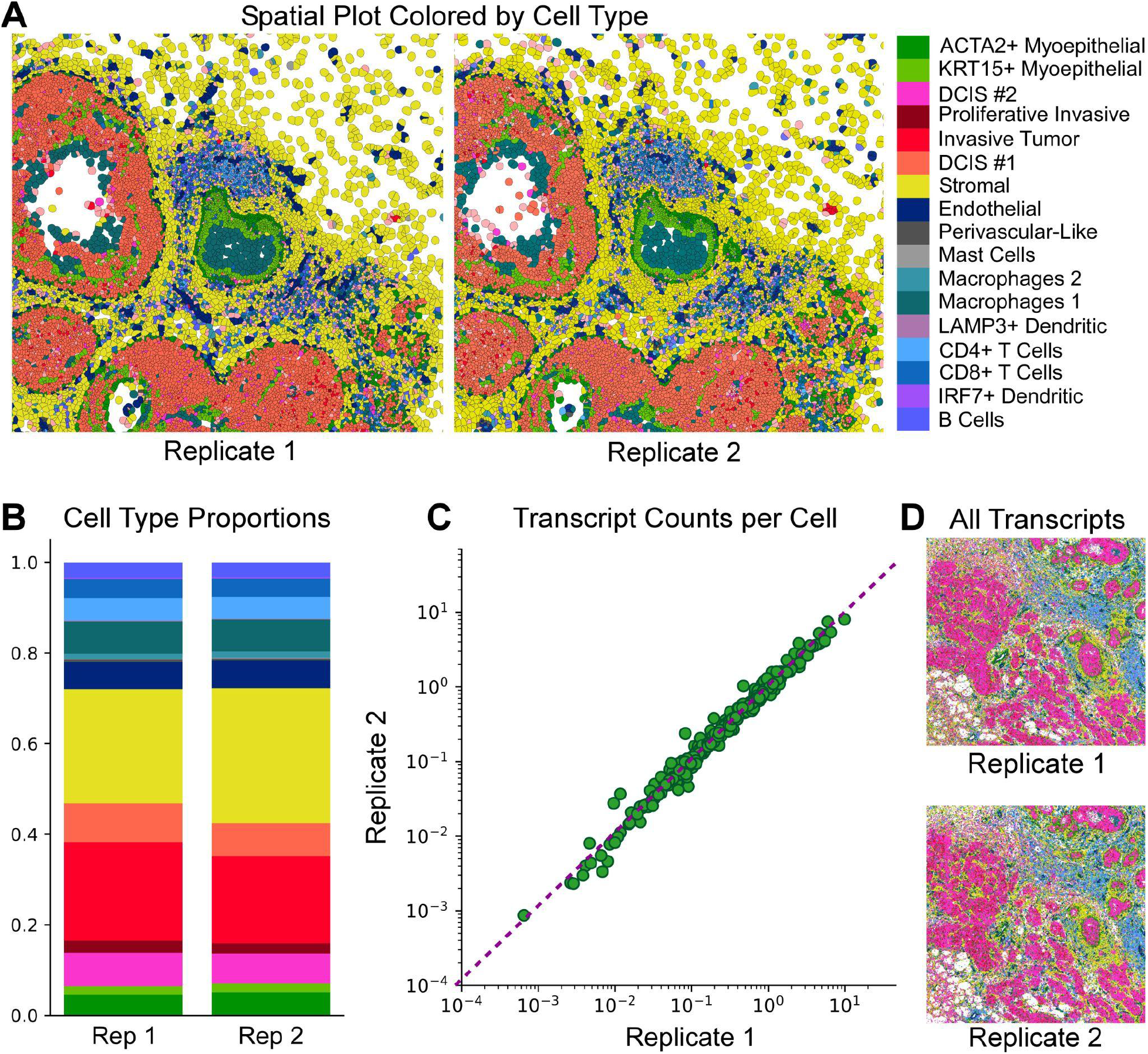
Serial section Xenium replicates are highly correlated. (A) Cell types for each replicate from the same ROI. (B) Cell type proportions across the whole section. (C) Scatter plot comparing transcripts per cell for each gene across replicates (r^2^ = 0.99). (D) ROI showing all transcripts, color coded by cell type, for each replicate.

**Supplemental Figure 7.**
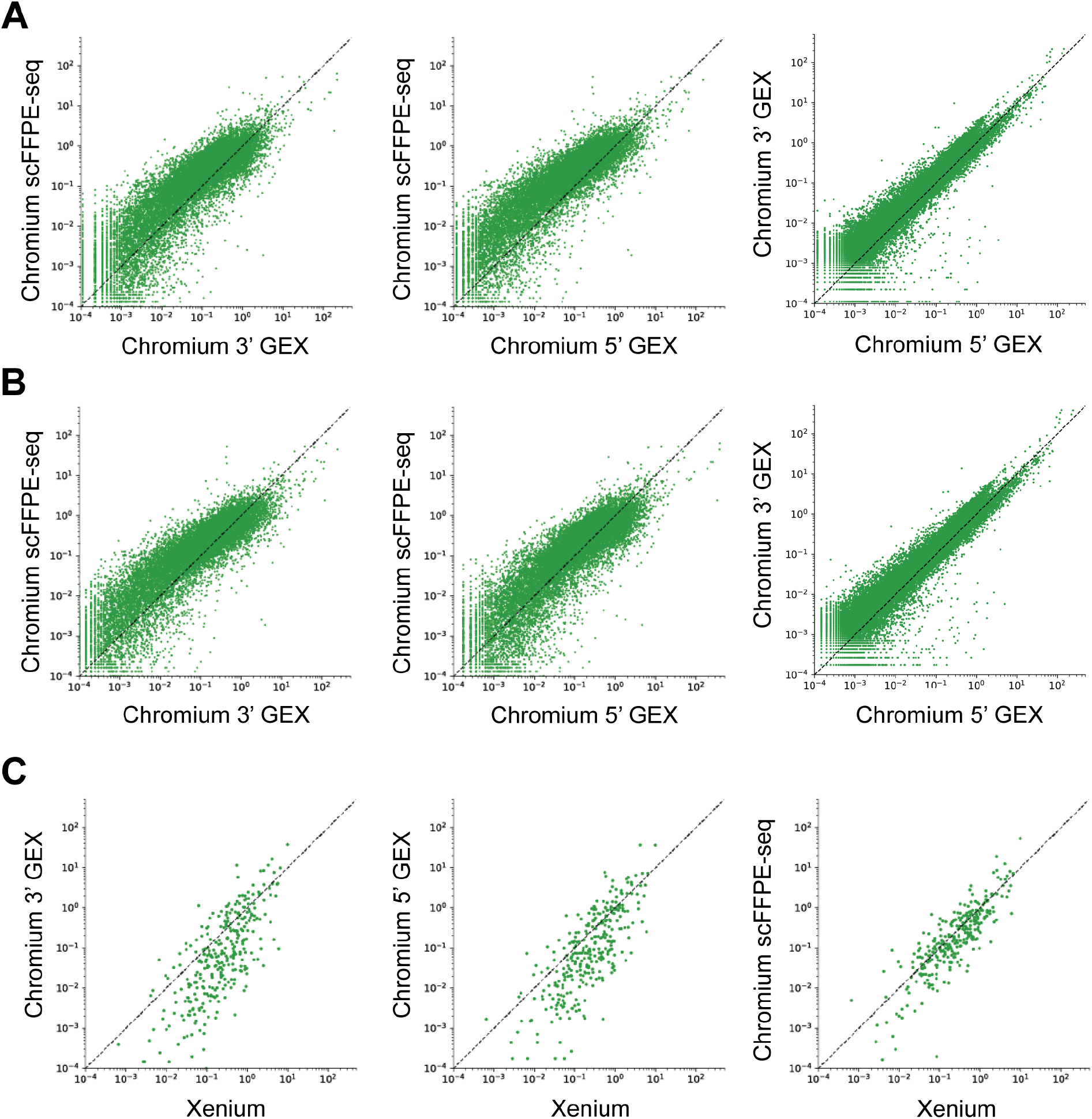
Benchmarking scFFPE-seq and Xenium sensitivity from FFPE curls against Chromium 3’ and 5′ GEX from patient-matched FF dissociated tumor cells. (A) Scatter plots of UMIs per cell representing pairwise comparisons of Chromium single cell technologies (scFFPE-seq, 3′ GEX and 5’ GEX) at the recommended sequencing depth for scFFPE-seq (10,000 mean reads per cell). (B) Scatter plots of UMIs per cell in pairwise comparisons between scFFPE-seq, 3′ GEX and 5’ GEX at the recommended sequencing depth for 3’ GEX and 5’ GEX (20,000 mean reads per cell). (C) Scatter plots of UMIs per cell and transcripts per cell between Chromium and Xenium. Genes are downsampled to those contained within the Xenium panel. Dotted lines represent X=Y.

**Supplemental Figure 8.**
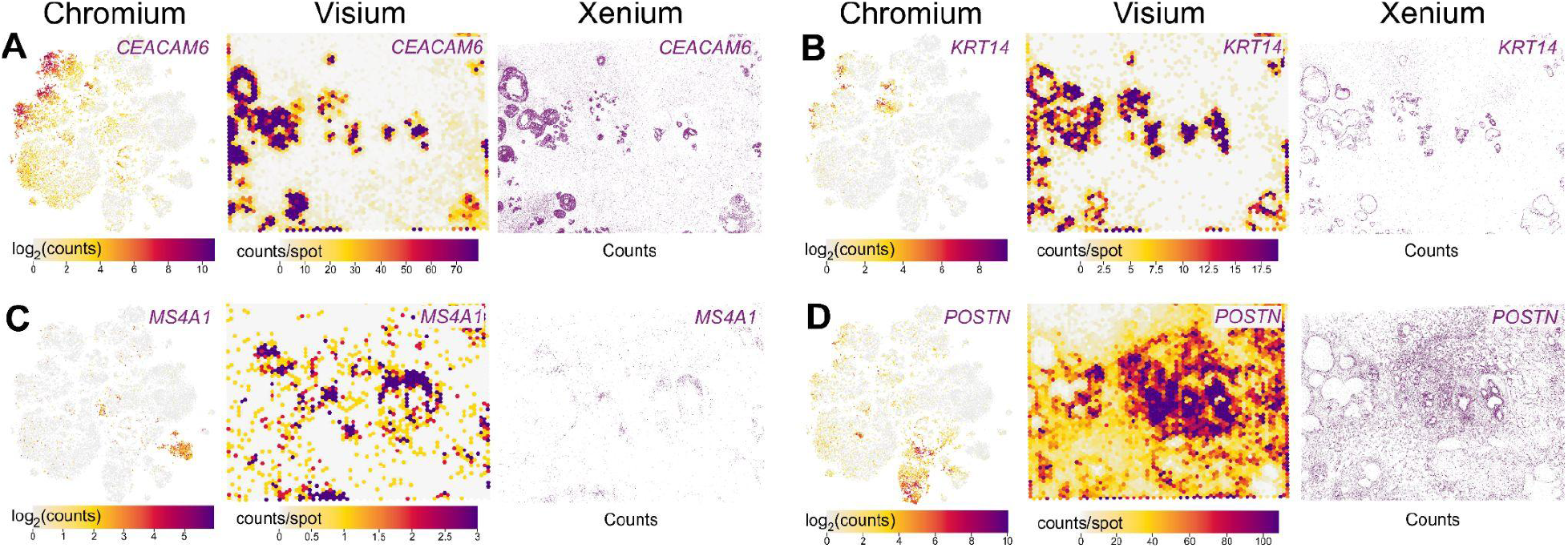
Comparison of Chromium, Visium, and Xenium FFPE technologies for key genes. A human breast cancer FFPE block was sectioned and processed for Chromium scFFPE-seq, Visium, and Xenium (see Fig. 1). (A) Human breast cancer oncogene *CEACAM6* is expressed in a subset of *ERBB2*+ (HER2+) tumor epithelial cells in the scFFPE-seq data. The Xenium spatial plot showing decoded transcripts reveals that *CEACAM6* is expressed exclusively in ductal carcinoma in situ (DCIS) regions, but is excluded from invasive cancer domains. Visium data validate this observation with high spatial correlation to the Xenium data. (B-D) Other exemplary markers for myoepithelial cells (*KRT14*), B cells (*MS4A1*), and stromal cells (*POSTN*) are shown across all three technologies, demonstrating cell type representation and spatial concordance.

**Supplemental Figure 9.**
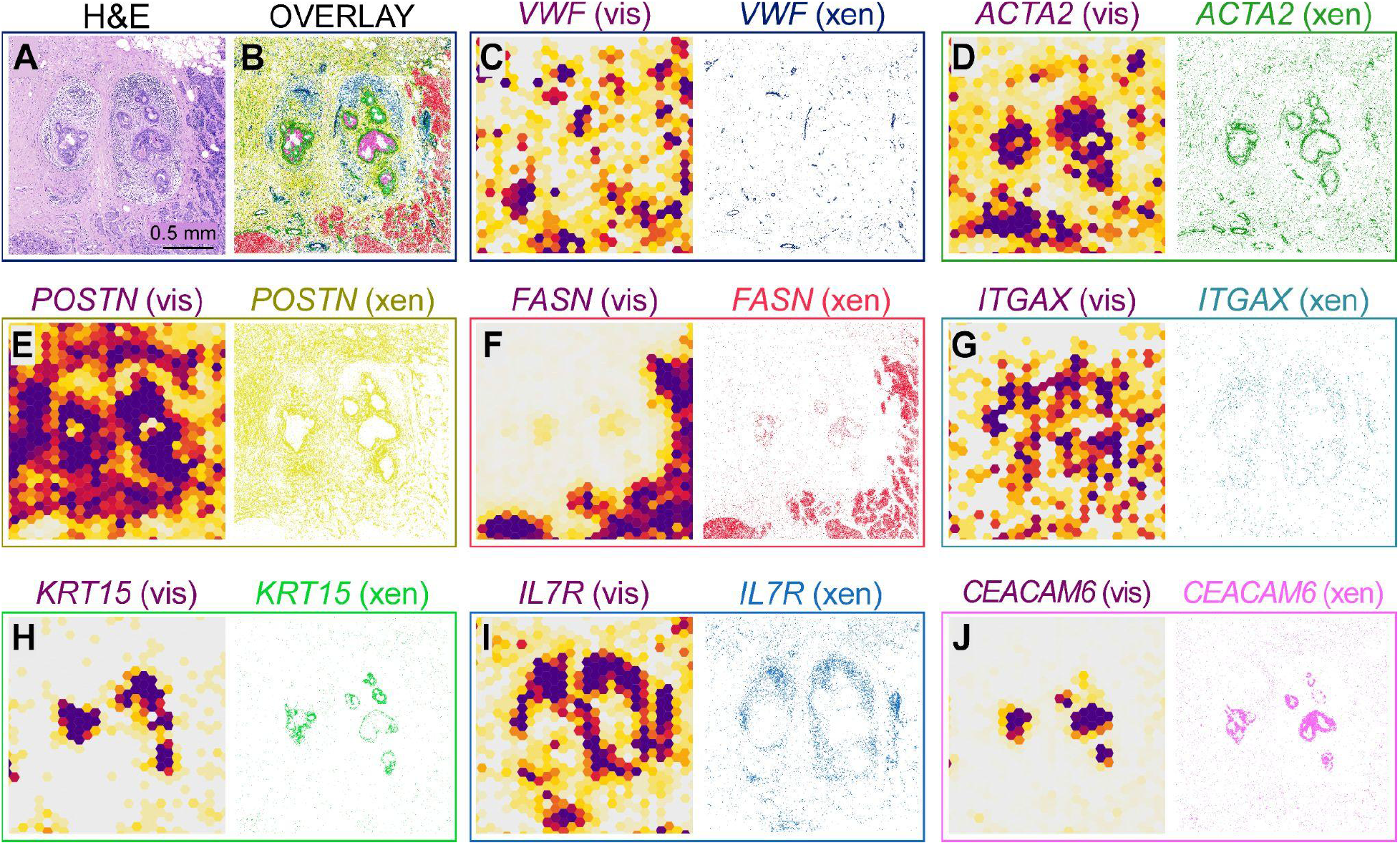
Comparison of Visium and Xenium for an ROI with high proximity and variety of cell types. Continuation of Fig. 4. (A) One small region of interest (ADH; atypical ductal hyperplasia) with a high diversity of cell types in close proximity. (B) Xenium spatial plot showing an overlay of eight selected gene markers. (C-J) All genes pictured in the overlay are shown individually for the same ROI with Visium (vis) on the left and Xenium (xen) on the right.

**Supplemental Figure 10.**
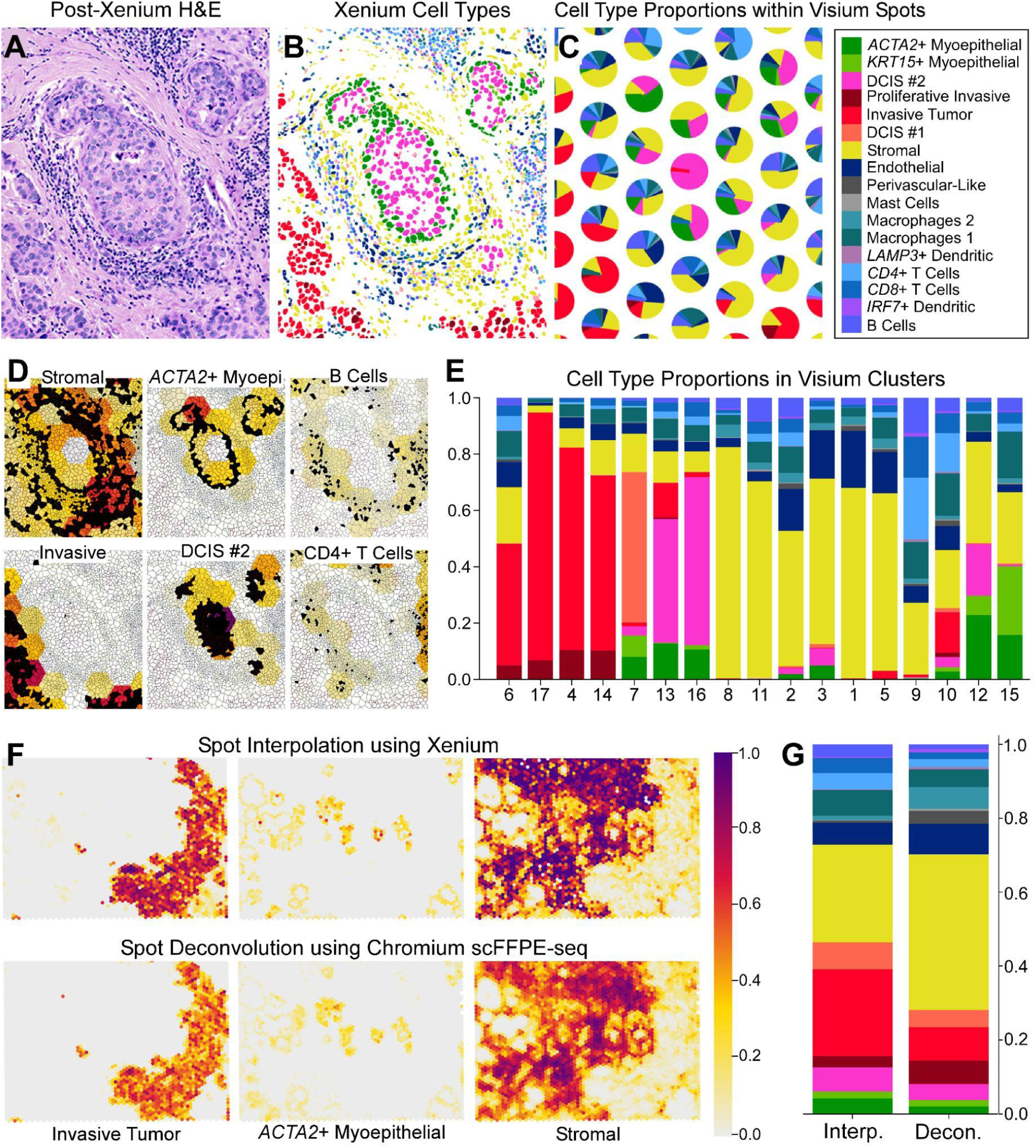
Xenium and scFFPE-seq assign cell type frequencies to individual Visium spots. (A-C) Registration of Visium and Xenium H&E images (see Fig. 4B) allows for the interpolation of cell type proportions within Visium spots using Xenium transcript counts. (D) Annotated cell types (see Supp. Fig. 3) from Xenium are overlaid on a Visium heatmap expressed as a proportion (fraction of one) of that cell type within each spot (see scale bar in F). (E) Interpolation of Visium spots allows for mixed, unannotated cell clusters from Fig. 3B to be resolved into cell type frequencies. (F) We deconvolved Visium spots into cell types with spacexr (Cable et al. 2021) using the scFFPE-seq data as the single cell reference. (F-H) Comparison of the spot interpolation method using Xenium and the deconvolution method using scFFPE-seq. Cell type proportions are expressed as a fraction of one.

### Supplemental Information

**Supplemental Table 1.** Metrics.

|  |  |  |  |  |  |  |  |  |  |  |  |  |  |  |
| --- | --- | --- | --- | --- | --- | --- | --- | --- | --- | --- | --- | --- | --- | --- |
| scFFPE-seq | Tissue (Curl) Size | Tissue Type and Thickness | Number of Cells | Total Transcripts | Median Reads per Cell | Median Genes per Cell | Genes Targeted | Genes Detected | Median UMI Counts per Cell | Reads Mapped Confidently to Probe Set | Pipeline Version | Fraction mitochondrial UMI counts | Estimated UMIs from Genomic DNA | Probe Set Name |
|  | 7.5 mm x 6 mm | 50 µm FFPE curls | 30,365 | 204,470,550 | 10,450 | 1,480 | 18,085 | 17,696 | 2,191 | 93.20% | cellranger-7.0.1 | 4.00% | 0.04% | Human Transcriptome Probe Set v2 |
| Visium | Tissue (Capture) Size | Tissue Type and Thickness | Number of Spots Under Tissue | Total Transcripts | Mean Reads per Spot | Median Genes per Spot | Genes Targeted | Genes Detected | Median UMI Counts per Spot | Reads Mapped Confidently to Probe Set | Pipeline Version | Fraction mitochondrial UMI counts | Estimated UMIs from Genomic DNA | Probe Set Name |
|  | 6.5 x 6.5 mm | 5 µm FFPE | 4,992 | 114,568,641 | 40,003 | 5,712 | 18,085 | 18,056 | 13,973 | 98.50% | spaceranger-2.0.0 | 6.60% | 3.00% | Human Transcriptome Probe Set v2 |
| Xenium | Tissue (Imaging) Size | Tissue Type and Thickness | Number of Cells | Transcripts Decoded (Q-score ≥ 20) | Median Reads per Cell | Median Genes per Cell | Genes Targeted | Genes + Add-On | Median Transcripts per Cell | Reads Mapped Confidently to Probe Set | Fraction Counts Neg. Control Probes | Fraction Counts Decoding Controls | Fraction Counts from Genomic DNA | Probe Set Name |
|  | 7.5 mm x 6 mm | 5 µm FFPE | 167,885 | 36,944,521 | NA | 62 | 313 | 280+33 | 166 | NA | 0.026% | 0.01% | 0.00% | Human Breast Panel |

**Supplemental Table 2.**
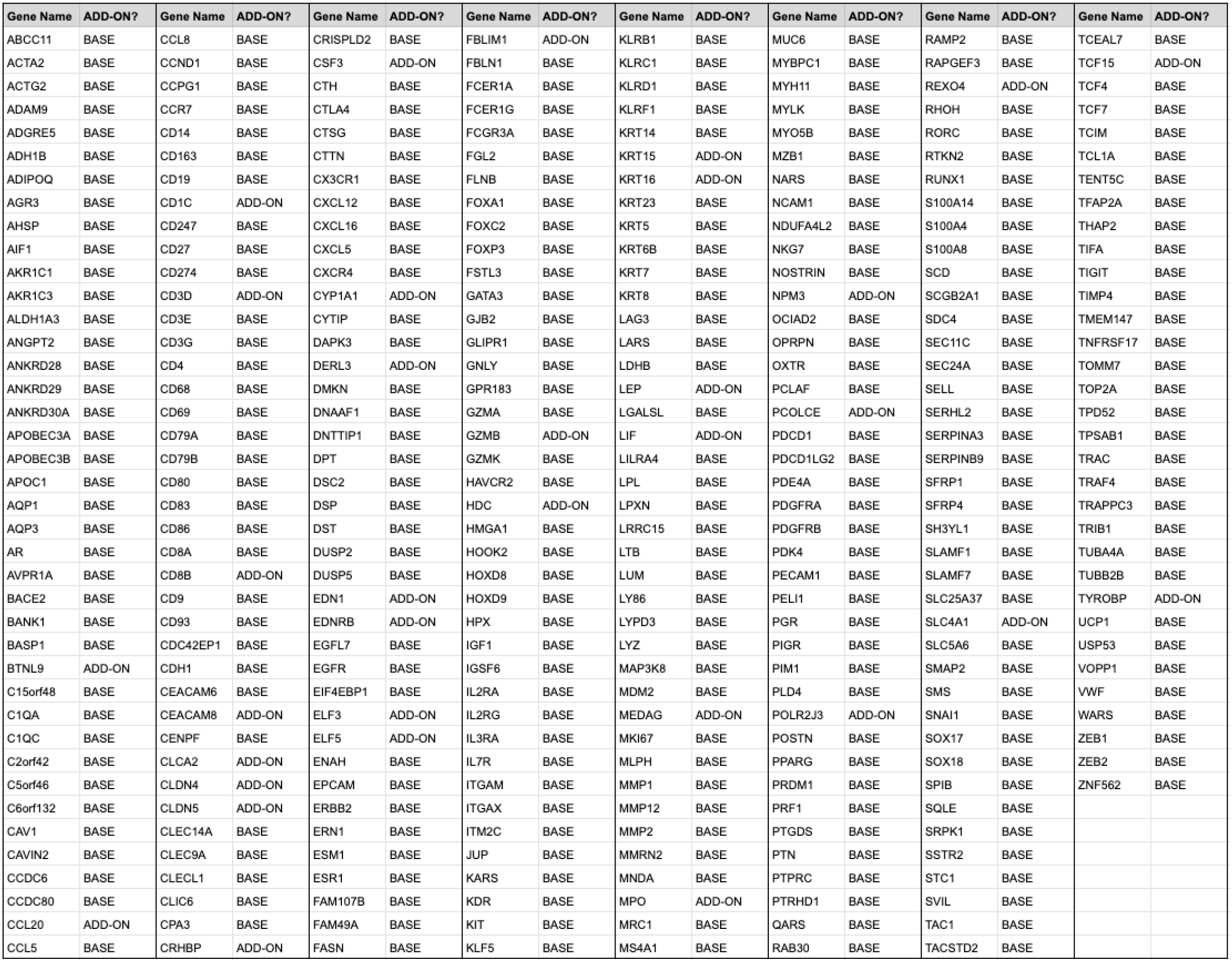
Xenium Gene List.

## 10x Development Teams

10x Genomics team members who all contributed to the development of technologies used in *High resolution mapping of the breast cancer tumor microenvironment using integrated single cell, spatial and in situ analysis of FFPE tissue*

Francis Aguisanda, Matt Alexander, Maria Alexis, Pratomo Alimsijah, Morris Allison, Aditya Aman, Joseph Aman, Zaid Ammari, Naishitha Anaparthy, Eric Anderson, Courtney Anderson, Melissa Ando, Joey Arthur, Adam Azarchs, Navi Bains, Naveed Bakh, Alexandru Bardasu, Carlos Barraza, Ashley Basco, David Batey, Florian Baumgartner, Alessio Bava, Kamila Belhocine, Jason Bell, Julio Beltran, Alex Ben, Zachary Bent, Casey (Jon) Berridge, Rajiv Bharadwaj, Anton Bjorninen, Kirk Blackmoore, Octavian Bloju, Lorita Boghospor, Jeff Bolland, Raj Borade, Erik Borgstrom, Sebastian Le Bras, Christian Broms, Phillip Brooks, Brenden Brown, Noel Harris Brown, Elizabeth Buckley, Patricia Buenbrazo, Ratha Bun, Diana Burkart-Waco, Jim Burrows, Michele Busby, Janae Bustos, Michele Caceres, Matthew Cai, Xiaoyang (Frank) Cai, Michael Campbell, Saranya Canchi, Joshua Cataldo, Emily Chan, Rena Chan, Alan Chang, Evelyn Chang, Ella Hsinwen Chang, Sharmila Chatterjee, Sidharth Chaturvedi, Mitu Chaudhary, James Chell, Kevin Chen, Sarah Chen, Hong Chen, Jui-Szu Chen, Sharon Chen, Brian Cheng, Jennifer Cheung, Jennifer Chew, Jun Ding Chiang, Hariharasudhan Chirra, Cheyenne Christopherson, Sonya Clark, Brynn Claypoole, Justin Costa, Julia Cowen, David Cox, Francis Cui, Abbey Cutchin, Aos Dabbagh, Smritee Dadhwal, Eileen Dalin, Aldo DeAmicis, Filip Defoort, Evan Dejarnette, Joshua Delaney, Nigel Delaney, Loruhama Delgado-Rivera, Maurizio Depoli, Abou Diop, Rhyan Dockter, Keri Dockter, Sultan Doganay, Jason Downing, Tyler Drake, Shoshoni Droz, Narek Dshkhunyan, Jens Durruthy Durruthy, Carine Edder, Peter Edge, Aaron Ellis, Emre Erhan, James Erhard, Cedric Espenel, Danielle Estes, Eric Evje, Navid Farahani, Reynaldo Farias, Dina Finan, Melanie Freeman, Danny Freitas, Tristan French, Meghan Frey, Alex Gagnon, Caroline Gallant, Abigail Gallegos, Christina Galonska, Karthik Ganapathy, Richard Gantt, Jason Gao, Kevin Gilmartin, Ryan Gilmore, Gabriele Girelli, Andreas Girgensohn, Michael Glazer, Shalini Gohil, Sumanth Gollapudi, Qiang Gong, Brandon Gordon, Erica Graziosa, Yang Gu, Josh Gu, Zhenping Guan, Zhengping Guan, Christopher Guido, Madhu Gundapuneni, Vaibhav Gupta, Rashmi Gupta, Lauren Gutgesell, Daniel Gyllborg, Lila Haba, Katherine Habeck, Andrew Haling, Katrín Halldórsdóttir, Andrej Hartnett, Ryo Hatori, Eileen He, Andrew Heiberg, Robert Henley, Janine Hensel, Lance Hepler, Alexander Hermes, Javier Hernandez, Iván Hernández, Jack Herrera, Jill Herschleb, Andrew Hill, Ben Hindson, Michael Hirte, Khoa Hoang, David Hoffman, Catie Hollyer, Eric Holt, Babak Honaryar, KC Hong, Daniel Horvath, Jennifer Hsu, Alice Huang, Winnie Hui, Kim Xuan Huynh, Hayato Ikoma, Moussa Iskandar, Eswar Iyer, Shaun Jackman, Sunil Jagannath, Samir Jain, Thomas Johnson, Tom Johnson, Guy Joseph, Aleksandra Jurek, Aarushi Kalaimani, Govinda Kamath, Lily Blanche Kameny, Ivar Karam, Rishabh Kasliwal, Layla Katiraee, Anna-Maria Katsori, Matthew Keeshen, Andrew Kessler, Tousif Khan, Aliyya Khan, Hanyoup Kim, Albert Kim, James Kim, Minji Kim, Sugyeom Kim, Alex Kindwall, Naga Sudha Kodavatikanti, Sutheng Kok, Sukhmal Kommidi, Nikola Kondov, Aishwarya Konnur, Jason Koth, Maria Kourbatov, Joseph Kovac, Sreenath Krishnan, Benjamin Ku, Jing Kuang, Malte Kuhnemund, Divya Anjan Kumar, Vijay Kumar, Josy Kuriakose, Sural Labha, Yves Lacroix, Karen Lai, Jana Lalakova, Tim Lam, Du Linh Lam, Alyssa Lanza, Jonathan Lau, Eunice Lau, Julia Lau, Lily Le, Soo Hee Lee, Thomas Lee, Josephine Lee, Junhyuck Lee, Elizabeth Lee, Jennifer Lew, Meryl Lewis, Peigeng Li, Vera Li, Ziyang Li, Qiang Li, Yuwei Li, Dongyao Li, Alvin Liang, Varoth Lilascharoen, Marisa Lim, David Little, Kendra Liu, Evelyn Lo, Ludmila Lokteva, James Longmire, Glory Lopez, Steve Losh, Han Lu, Grace Lucas, Mike Lucero, Susana Lau Lui, Paul Lund, Yi Luo, Chris Macklin, Diego Magdaleno, Amanda Mah, Shamoni Maheshwari, Amit Maheshwari, Sarah Mahmoud, Michelle Mak, Liza Man, Arec Manoukian, Francis Marcogliese, Hendricus Marindra, Pat Marks, Patrick Marks, Allison Martin, Kristen Martins-Taylor, Alicia McCarthy, Benjamin Mccreath, Shane McDermed, Jeff Mellen, Francesca Meschi, Paulius Mielinis, Marco Mignardi, Nabil Mikhaiel, Pradyumna Mishra, Adam Monkowski, Lysette Moreno, David Morgan, Logan Morrison, Fing Moua, Nima Mousavi, Aditi Mukherjee, Donald Mullane, Veronica Gonzalez Munoz, Laura Munteanu, Sivaram Muthusubramanian, Gambhir Nagaraju, Monica Nagendran, Jay Nagin, Sovann Nak, Akshay Nakra, Ashkan Beyranvand Nejad, Amber Nelson, Brenda Nguyen, Vu Nguyen, William Nitsch, Amy Oh, Jean-Francois Olivier, Mimmi Olofsson, Megan Olsen, Brett Olsen, James Ong, Susana Ordaz, Jessica Östlin, Dulce Ovando-Morales, Alex Palacio, Rushi Panchal, Janice Papartassee, Abhijna Parigi, Minkyung Park, Anuj Patel, Shyam Patel, Sid Patel, Kayne Patterson, Jeffrey Peck, Marissa Pennell, James Perna, Katherine Pfeiffer, Kristen Pham, Caio Porto, Denis Pristinski, Nicholas Provenzano, Benjamin Pruitt, Abhi Puthenveetil, Teddy De Puy, Xiaoyan Qian, Yufeng Qian, Ken Quah, Ailen Quero, Mohammad Rahimi, Selva Rajendran, Martino Ramella, Tina Ramirez, Jaya Ramrakhyani, Harjeet Randhawa, Nikhil Rao, Nicole Rapicavoli, Karl Rauta, Ann Renschler, Rudy Rico, Daniel Riordan, Elijah Roberts, Guillaume Robichaud, Anatalia Robles, Blas Rodrigues, Nancy Conejo Rodriguez, Mary Rogawski, Patrick Rolli, Michael Rose, Holly Ross, Chris Rouillard, Mary Rowgaski, Mustafa Rupawalla, Spontaneous Russell, Paul Ryvkin, Aprana Sahajan, Janani Sampathkumar, Marcus Sands, Poonam Sansanwal, Ace Santiago, Jerald Sapida, Anuj Sareen, Didem Sarikaya, Fahima Sarker, Hiroshi Sasaki, Martin Sauzade, Michael Schnall-Levin, Jason Schultz, Laurie Scott, Preyas Shah, Ankur Shah, Nathan Shapiro, Nate Shapiro, Shankar Shastry, Kai Shen, Christian Shi, Shang Shi, Lisa Shi, Shreyas Shivalkar, Steve Short, Joe Shuga, Anton Shutov, Sana Siddiqi, Eric Siegel, Aisling Sinclair, Hardeep Singh, Pal Singh, Ben Sisserman, Alex Skrynnyk, Carl Skuce, Peter Smibert, Adam Smiechowski, Benjamin Smyth, Kelvin Soong, Delia Soto, Mario de Souza, Rapolas Spalinskas, Susanne Spiegelberg, Nico Sponer, Niranjan Srinivas, Yogesh Srinivas, Jasper Staab, William Stanislaus, Jacob Stern, Marlon Stoeckius, Ryan Stott, Zachary Strike, Arjun Sugumar, David Sukovich, Ilse Sweldens, Kokchuan Tan, Qiaoqiao Tan, Sarah Tang, Weiyi (Lily) Tang, Adrian Tanner, Raghu Tayanna, Ryan Taylor, Nolan Teasdale-Schaf, Augusto Tentori, Jessica Terry, Joshua Thao, Michael Tierney, Meiliana Tjandra, Mckenzi Toh, Jeremy Tong, Joaquin Torres, Thien Trac, Khoi Tran, Christopher Tran, Joaquin Trosper-Torres, Andriy Tsupryk, Hank Tu, Mesruh Turkekul, Cedric Uytingco, Dino Valdecanas, Miriam Valencia, Ben Veire, Toon Verheyen, Rajiv Verma, Jan Vigar, Priya Rajendran Vishnu, Olga Vorobyova, Dagmar Walter, Amywenli Wang, Su Wang, Jun Wang, Dylan Webster, Jason Weis, Neil Weisenfeld, Kevin West, Tobias Wheeler, Dieter Wilk, Chris Wing, Christopher Wing, Evan Winget, George Withers, Tiffany Wong, Alex Wong, Philip Wright, Snow Wu, Kevin Wu, Sherry Wu, Fen Xie, Xiankun Xu, Shawn Yackly, RaviTeja Yarlagadda, Jennifer Yeager, Yifeng Yin, Frances Yun, Brett Zaborsky, Negin Zaraee, Ryan Zeng, Meng Zhang, Thanutra Zhang, Yiran Zhang, Fuying Zheng, Orchid Zhu, and Yihui Zhu

